# Loop extrusion by cohesin plays a key role in enhancer-activated gene expression during differentiation

**DOI:** 10.1101/2023.09.07.556660

**Authors:** Rosa J. Stolper, Felice H. Tsang, Emily Georgiades, Lars L.P. Hansen, Damien J. Downes, Caroline L. Harrold, Jim R. Hughes, Robert A. Beagrie, Benjamin Davies, Mira T. Kassouf, Douglas R. Higgs

**Affiliations:** MRC Weatherall Institute of Molecular Medicine, Radcliffe Department of Medicine, University of Oxford, John Radcliffe Hospital, Oxford, OX3 9DS, UK; Chinese Academy of Medical Sciences Oxford Institute, Nuffield Department of Medicine, University of Oxford, Old Road Campus, Oxford, OX3 7BN, UK; Wellcome Centre for Human Genetics, Nuffield Department of Medicine, University of Oxford, Old Road Campus, Oxford, OX3 7BN, UK; Francis Crick Institute, London, NW1 1AT, UK

## Abstract

Enhancers and their target promoters often come into close physical proximity when activated. This proximity may be explained by a variety of mechanisms; most recently via cohesin-mediated chromatin loop extrusion. Despite this compelling hypothesis, acute depletion of cohesin does not cause widespread changes in gene expression. We have tested the role of cohesin-mediated loop extrusion on gene expression at the mouse alpha-globin locus during erythropoiesis. Acute depletion of cohesin downregulates alpha-globin expression at early but not late stages of differentiation. When single or multiple CTCF sites are placed between the alpha-globin enhancers and promoters, alpha-gene expression is downregulated. Importantly, the orientation of the CTCF site plays a critical role, suggesting that within this activated domain, cohesin predominantly but not exclusively translocates from the enhancers to the promoters. We find that loop extrusion does play an important role in establishing enhancer-promoter proximity and consequent expression of inducible genes during differentiation.

## Introduction

It is now widely accepted that the process of loop extrusion, mediated by the cohesin complex and delimited by convergently-orientated CTCF binding sites, plays a key role in packaging chromatin into large (100s-1000s kb) topologically associating domains (TADs)(Sanborn, Rao et al. 2015, Fudenberg, Imakaev et al. 2016). It has been noted that TADs often include both the enhancers and promoters of specific, independent regulatory domains and are separate from other similarly organised regulatory units(Symmons, Uslu et al. 2014). Structural variants disrupting TADs often lead to abnormal patterns of gene expression(Nora, Lajoie et al. 2012, Lupianez, Kraft et al. 2015, Tsujimura, Klein et al. 2015, Lupianez, Spielmann et al. 2016). Throughout lineage-specification and differentiation, sub-TADs, corresponding to enhancer-promoter interactions, appear, and it has been proposed that translocation of cohesin may juxtapose these linearly separated regulatory elements thus playing a role in initiating transcriptional activation. Contrary to what would be expected if this was the case, in most current reports, although TADs largely disappear when cohesin is downregulated, gene expression is not affected to the extent predicted(Nora, Goloborodko et al. 2017, Rao, Huang et al. 2017, Schwarzer, Abdennur et al. 2017, Wutz, Várnai et al. 2017, de Wit and Nora 2023).

Here, we have set out to further explore the role of cohesin and loop extrusion in the regulation of mouse alpha-globin gene expression. The alpha-globin genes lie in a 165 kb TAD that is present in several cell types(Oudelaar, Beagrie et al. 2020), but during differentiation of erythroid cells a 70 kb sub-TAD appears, containing the five alpha-globin enhancers and their cognate promoters lying 30-50 kb downstream. During differentiation, largely convergently orientated CTCF sites flanking the sub-TAD come into close proximity based on chromosome conformation capture and super-resolution microscopy(Davies, Oudelaar et al. 2017, Brown, Roberts et al. 2018). The onset and continuation of gene expression and transcriptional bursting correlates with the increased physical proximity of the enhancers and promoters. These findings are consistent with the loop extrusion hypothesis and the alpha-globin model thus provides an opportunity to test this experimentally.

We have tested the role of loop extrusion in enhancer-activated alpha-globin expression in two ways. First, we depleted cohesin during erythroid differentiation and assessed its effect on alpha-globin expression. Then, we took advantage of an alternative way of compromising loop extrusion, by placing an ectopic CTCF site at the alpha-globin locus. Once cohesin is loaded, it continues to translocate until it reaches a boundary element that blocks further translocation, or the loop extrusion complex dissociates from the chromatin fibre. Although this process may be stalled by a variety of protein complexes and nuclear processes such as transcription, the 11-zinc finger protein CTCF is thought to be the most important block to loop extrusion. Many CTCF binding sites and transcriptionally active genes are found at TAD boundaries(Dixon, Selvaraj et al. 2012). Interestingly, the CTCF binding sites flanking TADs and sub-TADs are usually convergently-oriented(Rao, Huntley et al. 2014). This can be explained by a specific interaction between the N-terminus of CTCF and the Rad21-Stag subunits of cohesin which is thought to occur when translocation is stalled(Li, Haarhuis et al. 2020, Pugacheva, Kubo et al. 2020). Using the alpha-globin model, we have investigated the effect of inserting one or multiple well characterised CTCF binding sites between the alpha-globin enhancers and promoters in either orientation. This has allowed us to test the influence of CTCF sites, in either orientation, in blocking translocation and the impact of this on gene expression.

Although cohesin-mediated loop extrusion is a popular model to explain enhancer-promoter interactions, experimental evidence addressing this is not yet conclusive, particularly for interactions of less than 100kb(Rinzema, Sofiadis et al. 2022); and the relatively small effects on gene expression is unexpected(de Wit and Nora 2023). Although loop extrusion may explain how enhancers and promoters come into close proximity, other explanations have been proposed. Indeed, in certain instances cohesin seems to interrupt cis-interactions, thereby disrupting transcription and gene expression(Rinzema, Sofiadis et al. 2022). Here, by analysing a single locus that is progressively induced during differentiation, we show that cohesin-mediated loop extrusion provides a plausible explanation of how enhancers and promoters come into proximity during differentiation and that interrupting this process leads to significant changes in gene expression.

## Results

### Alpha-globin expression is sensitive to cohesin depletion at the earlier stages of erythroid differentiation

To test the importance of cohesin in the regulation of alpha-globin expression, we created a mouse ES cell line in which both copies of the cohesin subunit Rad21 are tagged with FKBP. This allows for rapid degradation of Rad21 using the dTAG system(Nabet, Roberts et al. 2018) and thereby depletion of the cohesin complex. We differentiated these cells along the erythroid lineage, using a well-established *in vitro* embryoid body (EB) differentiation method(Francis, Harold et al. 2022). During the EB differentiation, cells start to express globin genes around day 5 but their expression increases on subsequent days; previously, erythroid cells isolated from day 7 EBs were established as an optimal population to study alpha-globin expression(Francis, Harold et al. 2022) (Figure 1a).

**Figure 1.**
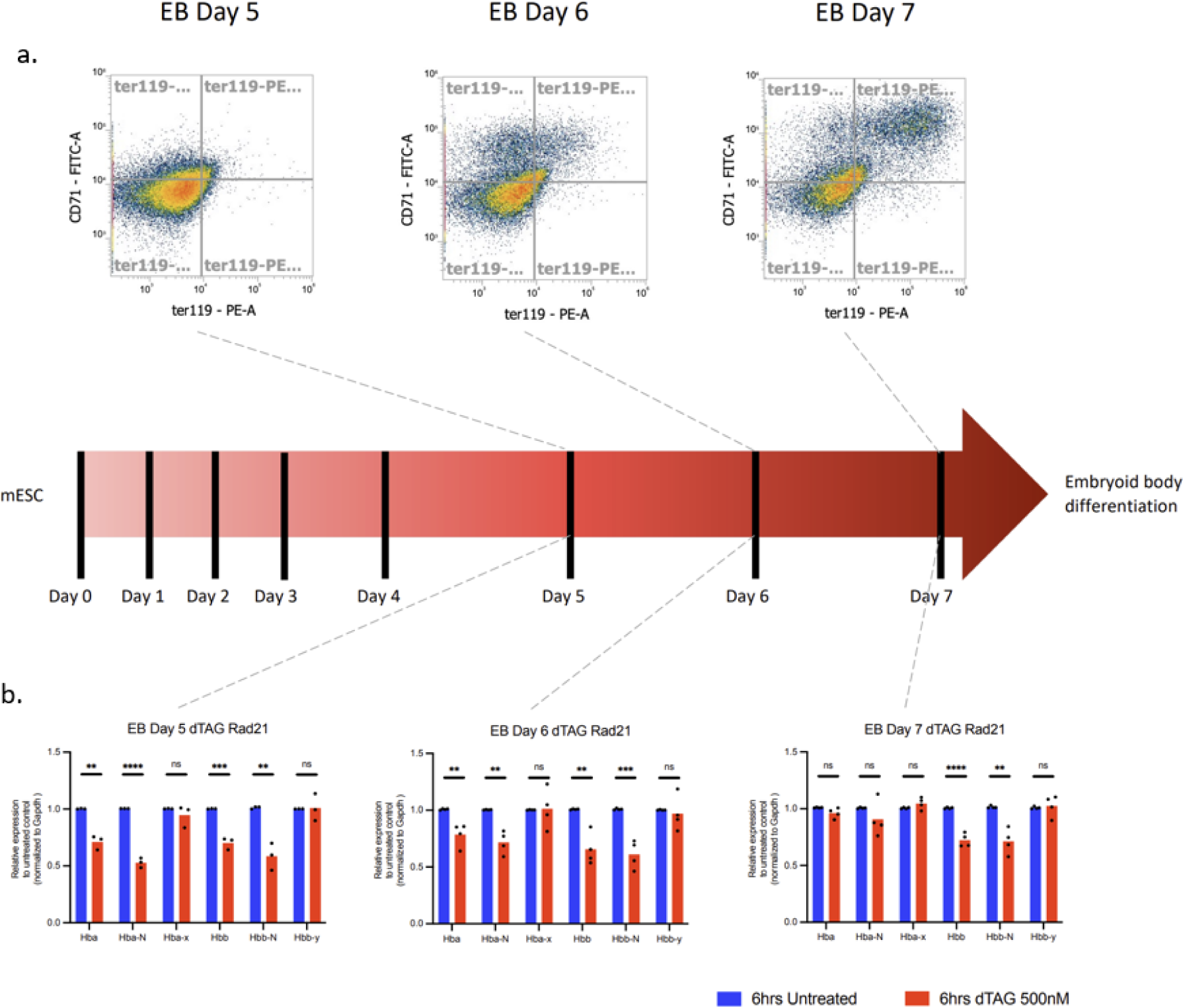
Alpha-globin expression is sensitive to cohesin depletion at the earlier stages of erythroid differentiation. a) During the 7-day differentiation protocol, mouse ES cells differentiate along the erythroid lineage and associated cell surface markers (CD71 and Ter119) increase. b) Gene expression as measured by qPCR normalised to Gapdh in day 5, day 6 and day 7 EB-derived erythroid cells with or without dTAG treatment. P-values were obtained by an unpaired two-tailed Student’s t-test: Not significant (NS) p > 0.05, * p <0.05, ** p < 0.01, *** p <0.001, **** p <0.0001.

Initially we purified erythroid Rad21-FKBP cells from day 7 EBs and treated this population with 500 nM dTAG for 6 hours, which led to rapid and complete degradation of Rad21 as judged by Western blot analysis (Supplementary figure 1). We determined expression levels of alpha- and beta-globin (*Hba-a* and *Hbb-b*) using RT-qPCR and observed no change in alpha-globin expression. However, we noted a 28% reduction of beta-globin expression (Figure 1b). We also used primers that amplify the intron-exon junctions of the alpha- and beta-globin genes and therefore give insight into nascent transcription, which shows similar patterns to those of mature transcripts. Finally, we looked at the embryonic globin genes (*Hba-x* and *Hbb-y*) and observed no change in their levels of expression following depletion of cohesin.

Expression of beta-globin is normally delayed relative to alpha-globin in the EB differentiation system(Francis, Harold et al. 2022). We wondered if this played a role in the sensitivity of beta-but not alpha-globin expression to cohesin degradation at day 7. Furthermore, of interest, it has previously been shown that cohesin may be especially important during changes in gene expression rather than in steady-state expression(Cuartero, Weiss et al. 2018). We therefore targeted earlier populations of erythroid cells, isolated from day 5 and 6 EBs, when expression is changing from very low to very high levels. Depleting Rad21 for 6 hours in these earlier populations had a clear and significant effect on both alpha- and beta-globin RNA expression (Figure 1b), with the less mature erythroid cells derived from day 5 EBs showing the most severe effect on alpha-globin expression upon cohesin depletion (29%). However, the expression of non-erythroid genes did not change (Supplementary figure 2).

By contrast to many previously analysed genes, expression of the globin genes, which is highly induced during the early stages of erythroid differentiation, is sensitive to the acute depletion of cohesin.

### Insertion of a CTCF insulator element in either orientation between the alpha-globin enhancers and the alpha-genes

Translocation of cohesin can be blocked by CTCF, and we therefore reasoned that placing a CTCF binding site ectopically between the alpha-globin enhancers and the alpha-globin genes would be an orthogonal way of testing the importance of loop extrusion for alpha-globin gene activation. The insertion of an ectopic binding site would offer the benefit of blocking loop extrusion throughout differentiation in a locus-specific manner. Additionally, since CTCF predominantly but not exclusively blocks loop extrusion from a specific direction in relation to its non-palindromic binding site, inserting it in either orientation could offer insight into the direction of loop extrusion. We therefore aimed to create two models: one with an ectopic CTCF binding site oriented towards the alpha-globin enhancers, blocking loop extrusion from that direction, and one in which the binding site is oriented to block loop extrusion from the direction of the promoters.

To select a CTCF binding site sequence to ectopically insert, we considered the CTCF binding sites within the alpha-globin locus that have been mutated and characterised in detail in previous studies(Hanssen, Kassouf et al. 2017, Harrold, Gosden et al. 2020). Although deletion of these CTCF sites led to changes in chromatin interactions, deletion of nearly all single CTCF sites had little or no effect on gene expression. The one exception was the deletion of a CTCF site (HS-38) located at the 5’ boundary of the alpha-globin sub-TAD(Oudelaar, Beagrie et al. 2020) (Figure 2a and b). This CTCF binding site normally restricts the influence of the alpha-globin enhancers on genes lying 5’ (upstream) of the globin cluster. Deletion of HS-38 leads to upregulation of these genes(Hanssen, Kassouf et al. 2017). DNaseI hypersensitivity data reveals two footprints for this binding site: one overlapping with the core motif for CTCF and one corresponding to a previously described upstream motif (Figure 2b)(Nakahashi, Kwon et al. 2013). This means it is possible to confidently determine the orientation of this binding site. Therefore, this binding site appeared a good candidate to block loop extrusion between the alpha-globin enhancers and the alpha-genes. We selected an 83-bp sequence, encompassing the entire HS-38 DNaseI footprint, and placed this sequence between the most 3’ enhancer (R4) and the most 5’ globin gene, *Hba-x* (Figure 2c).

**Figure 2.**
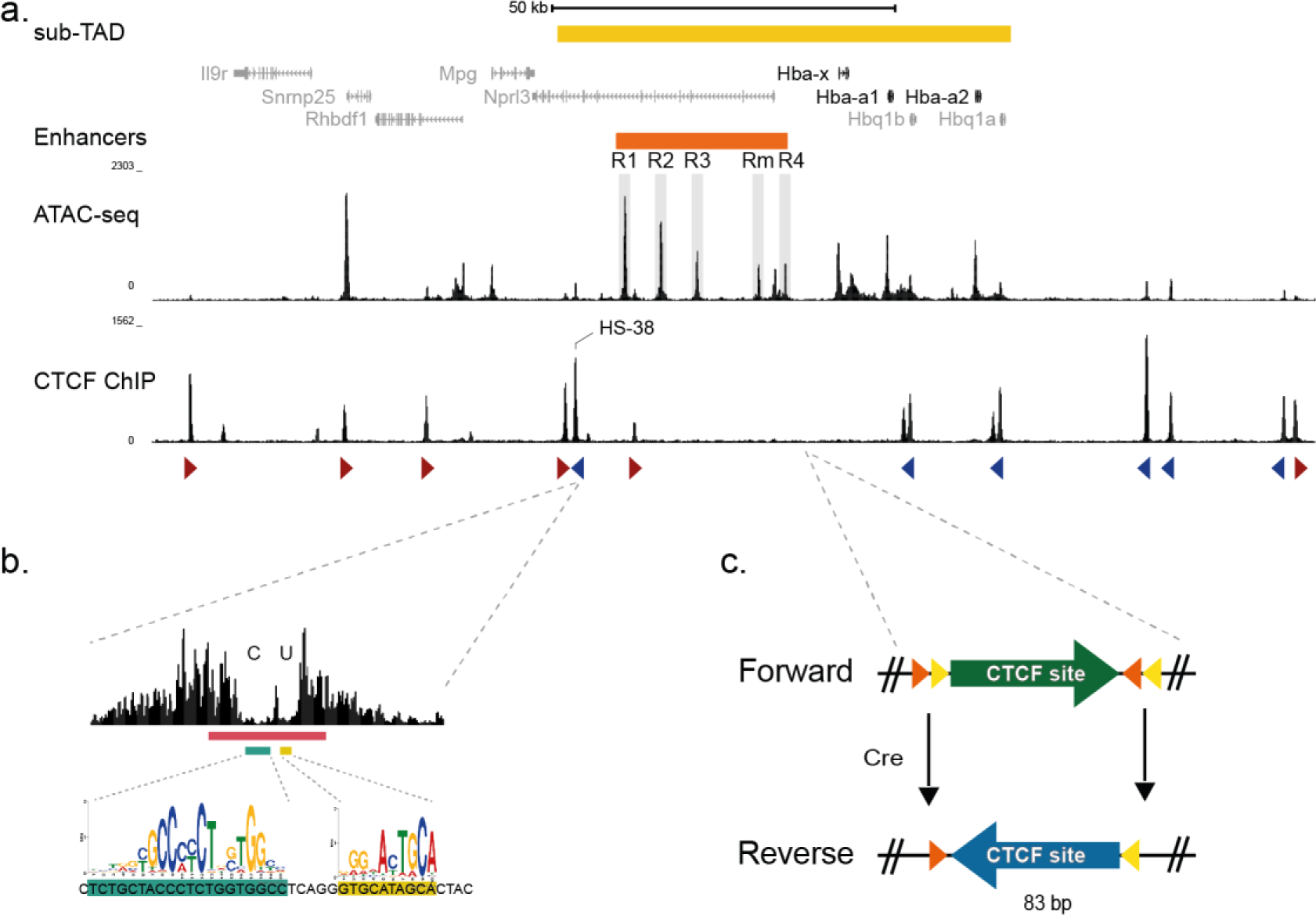
Insertion of a CTCF insulator element in between the alpha-globin enhancers and the alpha-genes. a) The mouse alpha-globin locus lies in a 65 kb sub-TAD (yellow bar) and contains two adult alpha-globin genes (Hba-a1 and Hba-a2) and an embryonic gene, zeta-globin (Hba-x). Open chromatin peaks (ATAC-seq track(Francis, Harold et al. 2022)) mark five upstream enhancer elements, four of which are located in the introns of the Nprl3 gene. Several CTCF binding sites (CTCF-seq track(Francis, Harold et al. 2022)) flank the sub-TAD. b) CTCF binding site HS-38 has been previously identified as a boundary element(Hanssen, Kassouf et al. 2017). Its 50 bp DNaseI footprint shows a clear orientation with a core (marked C, green bar) and upstream (marked U, yellow bar) motif. The red bar indicates the 83 bp sequence used in this study. c) The 83 bp sequence indicated in b) was introduced between the alpha-globin genes and its enhancers, 5 kb upstream of the zeta-globin promoter. Heterospecific lox sites (orange triangle: lox2272, yellow triangle: loxP) flank the new binding site and enable the inversion of the binding site orientation after addition of Cre recombinase.

The CTCF site was initially oriented convergent with downstream CTCF binding sites and towards the alpha-globin genes, and we termed this the “Forward” model. The ectopic CTCF binding site was flanked by heterospecific lox sites, to enable its inversion to create the “Reverse” model, in which the CTCF site is convergent with upstream CTCF binding sites and oriented towards the enhancer cluster (Figure 2a and c). Initially, we used CRISPR/Cas9- mediated homology-directed repair to introduce this sequence into mouse ES cells, and following blastocyst injection, we also obtained a Forward and Reverse mouse model. Again, we also used *in vitro* EB differentiation to produce erythroid cells from the mouse ES cell models, using the day 7 timepoint. From the engineered mouse models, we analysed spleen and fetal liver-derived erythroid cells as well as E10.5 embryonic blood.

To confirm that CTCF binds to the newly introduced ectopic site, we generated CTCF ChIP-seq in both the Forward and Reverse models in EB- and fetal liver-derived erythroid cells (Figure 3). This verified CTCF binding to the newly introduced ectopic site; there were little, if any, changes at existing CTCF binding sites. Importantly, the ectopic ChIP-seq peak had a similar level of binding regardless of its orientation, showing efficient recruitment in both models. We also assessed the open chromatin landscape by generating ATAC-seq datasets in both wild-type cells and those with the ectopic insertion. Again, insertion of the ectopic CTCF binding site did not result in significant changes to surrounding hypersensitive sites compared to wild-type cells, establishing that the insertion does not disturb the overall integrity of the locus and the enhancers are still accessible (Supplementary figure 3). The new CTCF site is marked by a small accessible chromatin peak, similar to other CTCF binding sites at the locus.

**Figure 3.**
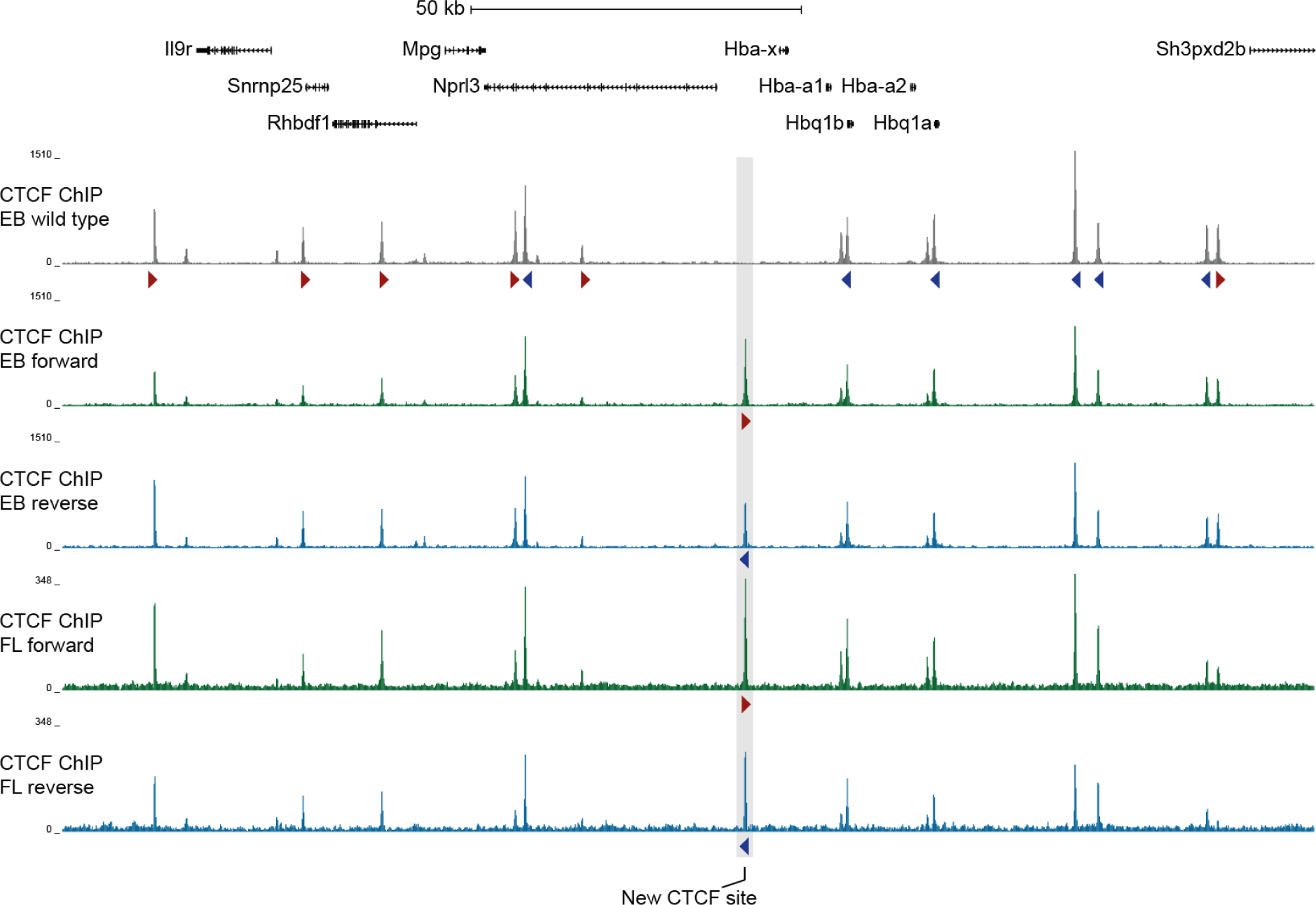
CTCF ChIP-seq confirms recruitment of CTCF to the newly introduced binding site. RPKM-normalised CTCF ChIP-seq tracks in wild-type(Francis, Harold et al. 2022) (grey), Forward (green) and Reverse (blue) day 7 EB-derived erythroid cells (EB, n=2 for each) and Forward and Reverse fetal liver-derived erythroid cells (FL, n=3 for each). CTCF binding site orientation is indicated by forward arrows (red) and reverse arrows (blue). The insertion site is highlighted in grey.

### Cohesin is present primarily at the reverse oriented CTCF site

Cohesin is usually present at most of the CTCF sites at the alpha-globin locus, and at a lower level at some other sites, such as the enhancers(Hanssen, Kassouf et al. 2017). To probe any changes in cohesin binding upon the introduction of the ectopic binding site, we performed ChIP-seq for cohesin subunit Rad21 in the Forward and Reverse model.

In EB-derived erythroid cells, there was a small peak of cohesin at the ectopic site in the Forward model (Figure 4a). In contrast, in the Reverse model there is a very prominent peak at the ectopic site. We observed the same pattern in spleen-derived erythroid cells, with a prominent peak in the Reverse model and a smaller peak in the Forward model (Figure 4a). This was surprising, as there was very little difference in CTCF recruitment between the Forward and Reverse model (Figure 3).

**Figure 4.**
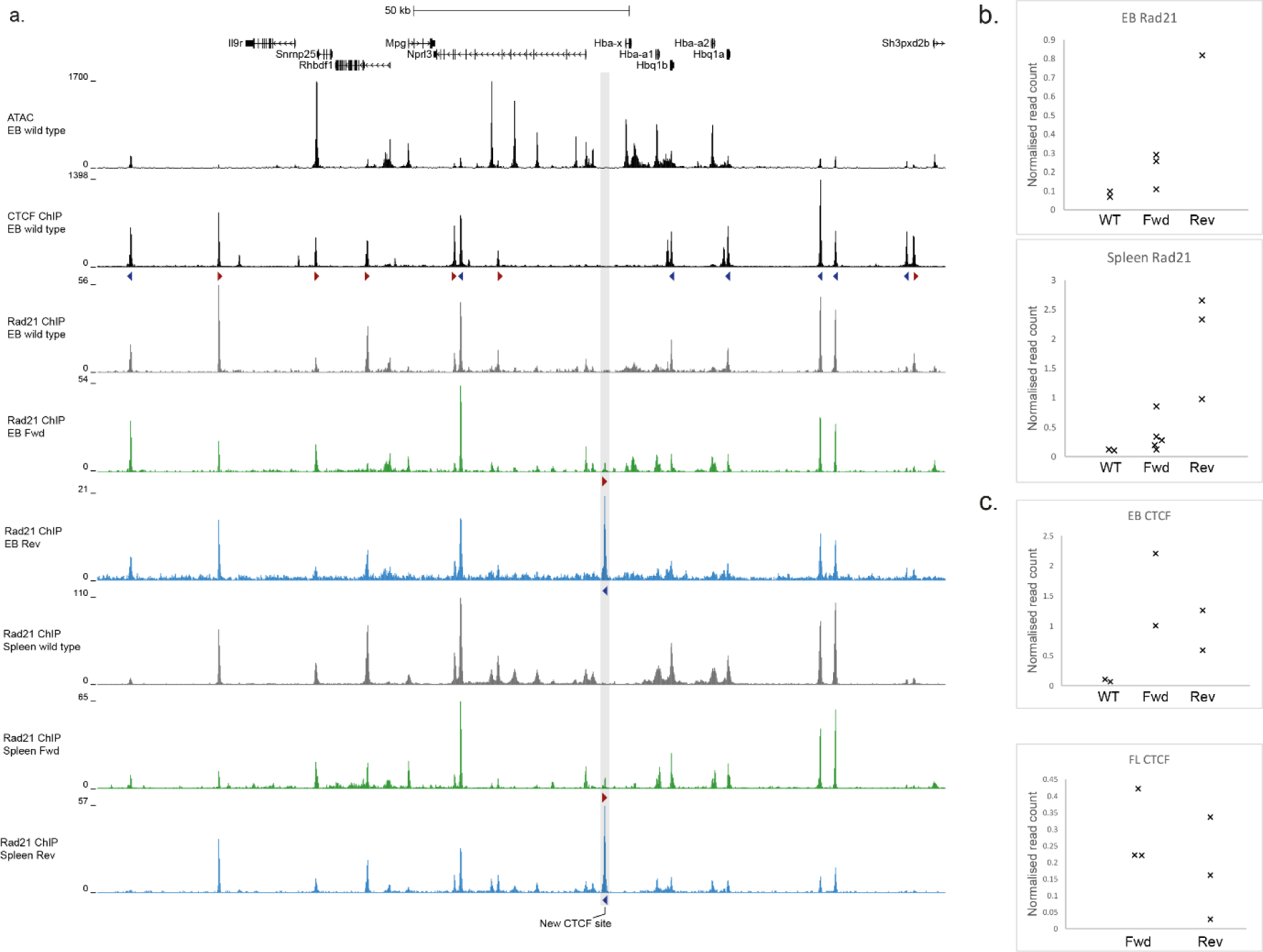
Rad21 ChIP-seq at the insertion site. a) Top two tracks: ATAC-seq and CTCF ChIP-seq tracks in wild-type EB-derived erythroid cells(Francis, Harold et al. 2022). RPKM-normalised Rad21 ChIP-seq tracks in wild type (grey, n=2), Forward (green, n=3) and Reverse (blue, n=1) in EB-derived erythroid cells, and below for spleen wild type(Hanssen, Kassouf et al. 2017) (grey, n=2), Forward (green, n=5) and Reverse (blue, n=3). CTCF binding site orientation is indicated by forward arrows (red) and reverse arrows (blue). The insertion site is highlighted in grey. b) Rad21 ChIP read counts over the insertion site in individual ChIPs, normalised to the average number of reads over all other peaks in the genome. c) CTCF ChIP read counts over the insertion site in individual ChIPs, normalised to the average number of reads over all other peaks in the genome.

We quantified the number of reads in a 1.5 kb window surrounding the ectopic CTCF binding site and normalised this to the average number of reads over genome-wide CTCF peaks (Figure 4b and c). This revealed that there is an enrichment of Rad21 at the ectopic site in the Forward model, but there are more reads in the Reverse model. In contrast, we observed that although the CTCF levels are variable, it is consistently present in both the Forward and Reverse model (Figure 4c).

Together, these findings suggest that while CTCF sites in either orientation affect the translocation of cohesin, the reverse orientation causes a more significant barrier to translocation than the forward orientation.

### CTCF sites oriented towards the enhancers cause the strongest reduction in alpha-globin expression

To assess if there is a difference in functional outcome between the Forward and Reverse model, we determined alpha-globin expression levels in various erythroid tissues using RT-qPCR. In EB-derived erythroid cells, alpha-globin expression was strongly reduced; expression was 44% lower in the Forward model compared to wild type, and 86% lower in the Reverse model (Figure 5a). We also determined levels of alpha-globin expression in E10.5 primitive embryonic erythroid cells, as these are the primary-tissue equivalents of EB-derived erythroid cells(Francis, Harold et al. 2022). In this tissue, alpha-globin expression is not significantly altered in the Forward model, but in the Reverse model expression is reduced by 68% (Figure 5a).

**Figure 5.**
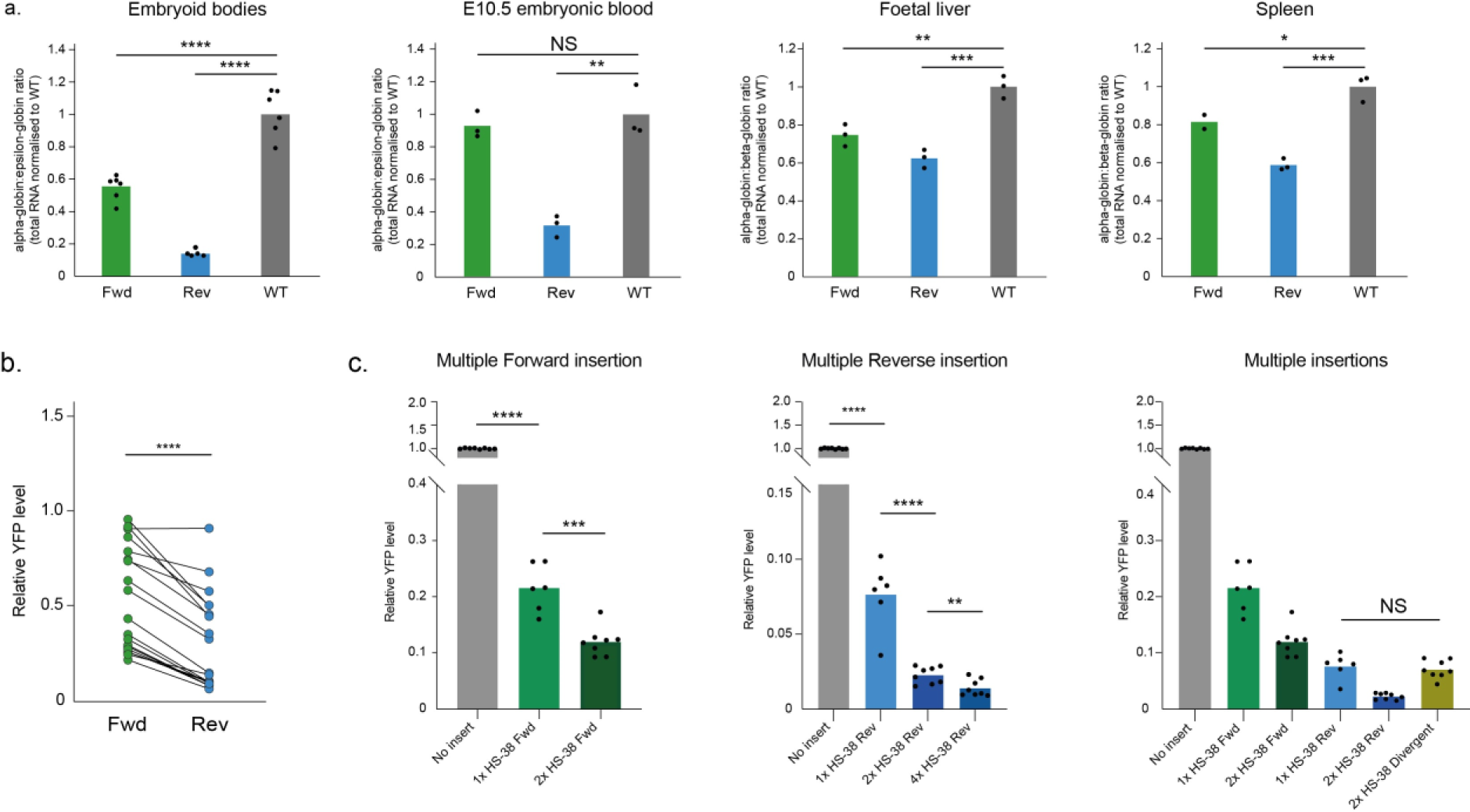
Expression of alpha-globin in various insertion models and erythroid tissues. a) Expression of alpha-globin in EB-derived erythroid cells, normalised to the expression of embryonic beta-like globin, epsilon-globin (Hbb-y; n=5 for Rev, n=6 for Fwd and WT), in E10.5 embryonic blood normalised to epsilon-globin (n=3 for all), in cultured E12.5 fetal liver-derived Ter119+ erythroid cells normalised to adult beta-globin (n=3 for all), and in spleen-derived Ter119+ erythroid cells normalised to beta-globin (n=2 for Fwd, n=3 for Rev and WT). Bar plot shows the mean ratio with individual data points marked by black dots. P-values were obtained using an unpaired two-tailed Student’s t-test. b) Relative YFP levels normalised to neutral insert control in clones with different CTCF binding site, either in forward or reverse orientation. P-value was obtained using a paired t-test. c) Relative YFP levels normalised to no insert control in clones with multiple CTCF insertions. P-values were obtained using an unpaired two-tailed Student’s t-test. Not significant (NS) p > 0.05, * p <0.05, ** p < 0.01, *** p <0.001, **** p <0.0001.

In two definitive, more adult-like, erythroid tissues, namely fetal liver-derived and spleen-derived erythroid cells, the alpha-globin expression is reduced both in the Forward and Reverse model. Again, the reverse orientation had the strongest effect on alpha-globin expression, with a 38% and 41% reduction respectively (Figure 5a).

Furthermore, we noted that the ectopic site did not change the expression of zeta-globin (*Hba-x*; Supplementary figure 4), much like Rad21 depletion had not (Figure 1b). This is in line with our previous observation that the expression of zeta-globin is predominantly independent of the alpha-globin enhancers(King, Songdej et al. 2021). However, surprisingly, the expression of several non-erythroid genes that lie outside of the sub-TAD but still within the larger (165kb) TAD was changed (Supplementary figure 5). Interestingly, the genes that increased in expression upon CTCF insertion, *Rhbdf1* and *Mpg,* were also affected when 5’ boundary CTCF sites were previously deleted(Hanssen, Kassouf et al. 2017). Thus, the ectopic HS-38 CTCF seems to not only affect alpha-globin enhancer-promoter interactions, but also disrupts the sub-TAD structure in general.

Having tested alpha-globin expression in multiple CTCF insertion models and several different erythroid tissues, a striking difference between the forward and reverse models emerged. In all erythroid tissues tested, the Reverse model, in which the CTCF site predominantly blocks cohesin translocating from the enhancers, had a stronger effect on alpha-globin expression. We wondered if this was a feature of this specific CTCF binding site, or of its position within the alpha-globin locus.

In a separate study(Tsang, Stolper et al. 2023), a reporter assay has been developed based on the observation made here: that alpha-globin expression is highly sensitive to disruption by a single CTCF binding site (Figure 5a). The reporter assay uses a YFP-tag at the alpha-gene to facilitate efficient evaluation of expression levels by FACS(Francis, Harold et al. 2022). This assay showed a range of alpha-globin expression when different CTCF sites were inserted(Tsang, Stolper et al. 2023).

As the sites were introduced both in the forward and reverse orientation, the data allowed us to study the contribution of binding site orientation further. We observed the same trend as in the Forward and Reverse models described here: a CTCF site orientated towards the enhancers forms a stronger insulator than when it is orientated towards the genes (Figure 5b). This shows that the orientation effect is not specific to the HS-38 binding site but is a general characteristic of the locus and the position of the CTCF sites relative to the enhancers and promoters.

### Insertion of multiple CTCF sites between the alpha-globin enhancers and the alpha- genes causes a further reduction in alpha-globin expression

Introduction of a single ectopic CTCF binding site, which blocks loop extrusion between the alpha-globin enhancers and genes, causes a strong reduction in alpha-globin expression, especially in cells derived from day 7 EBs. However, about 14% of alpha-globin expression remains in the Reverse model, and this is a substantial level of expression as alpha-globin is very highly expressed in wild-type erythroid cells. We therefore wondered if this reflected the limit of the contribution of loop extrusion to alpha-globin expression, or if it would be possible to reduce expression even further. It has been shown that a single CTCF binding site does not even block half of the translocating cohesin complexes(Davidson, Barth et al. 2023) and that additional CTCF sites can form a stronger boundary(Huang, Zhu et al. 2021).

We again employed the recently developed reporter assay(Tsang, Stolper et al. 2023) to test if we could disrupt alpha-globin expression even further. We designed several constructs, all inserting multiple copies of the same HS-38 CTCF binding site, spaced 500 bp apart. When two CTCF binding sites in the forward orientation were inserted, the insulation was strengthened, illustrated by a lower YFP level compared to one forward CTCF binding site (Figure 5c). Similarly, two reverse-oriented binding sites reduced the level of expression. Although expression levels are already low in cells with two reverse-oriented binding sites, adding two more to produce an allele with four reverse CTCF binding sites significantly lowers alpha-globin expression further still (Figure 5c). Importantly, introducing two CTCF binding sites in a divergent orientation (Figure 5c) did not increase the insulation compared to a single CTCF site in the reverse orientation.

### 3D chromatin architecture changes upon CTCF insertion in an orientation-dependent manner

The presence of Rad21 at the ectopic CTCF binding site seems to indicate that this newly inserted site alters loop extrusion and thus 3D genome organisation. However, it has also been shown that CTCF and cohesin localisation is very similar at CTCF binding sites between the undifferentiated, inactive alpha-globin locus, and the erythroid, active alpha-globin locus(Hanssen, Kassouf et al. 2017, Hua, Badat et al. 2021). Therefore, the 3D chromatin structure cannot be inferred from just the localisation of cohesin subunits and CTCF, and probing the 3D chromatin organisation can provide more information on how altering the translocation of cohesin may alter gene regulation at this locus.

We have previously shown by Capture-C(Davies, Telenius et al. 2016, Oudelaar, Beagrie et al. 2020), MCC(Hua, Badat et al. 2021) and by super-resolution imaging(Brown, Roberts et al. 2018) that the alpha-globin enhancers and promoters come into close juxtaposition within the erythroid specific sub-TAD both in primitive and definitive erythroid cells. As the sub-TAD forms, the convergent, flanking CTCF-elements are drawn into close proximity to each other. However, the flanking CTCF sites interact less frequently with the centre of the sub-TAD, thereby forming a loop structure encompassing the sub-TAD(Hanssen, Kassouf et al. 2017).

Using Capture-C, we investigated changes in the chromatin organisation upon introduction of the ectopic CTCF site in EB-derived erythroid cells. When the ectopic site was placed between the enhancers and promoters, in either orientation, interactions between these elements were reduced, when viewed from the main enhancer element R2 (Figure 6a)(Hay, Hughes et al. 2016). Of particular interest, there was a stronger reduction in the frequency of enhancer-promoter interactions when the ectopic CTCF site was orientated towards the enhancers (Reverse model; Figure 6a), where it should predominantly stall cohesin translocating from the enhancers to the alpha-globin genes. This is consistent with the changes we see in gene expression.

**Figure 6.**
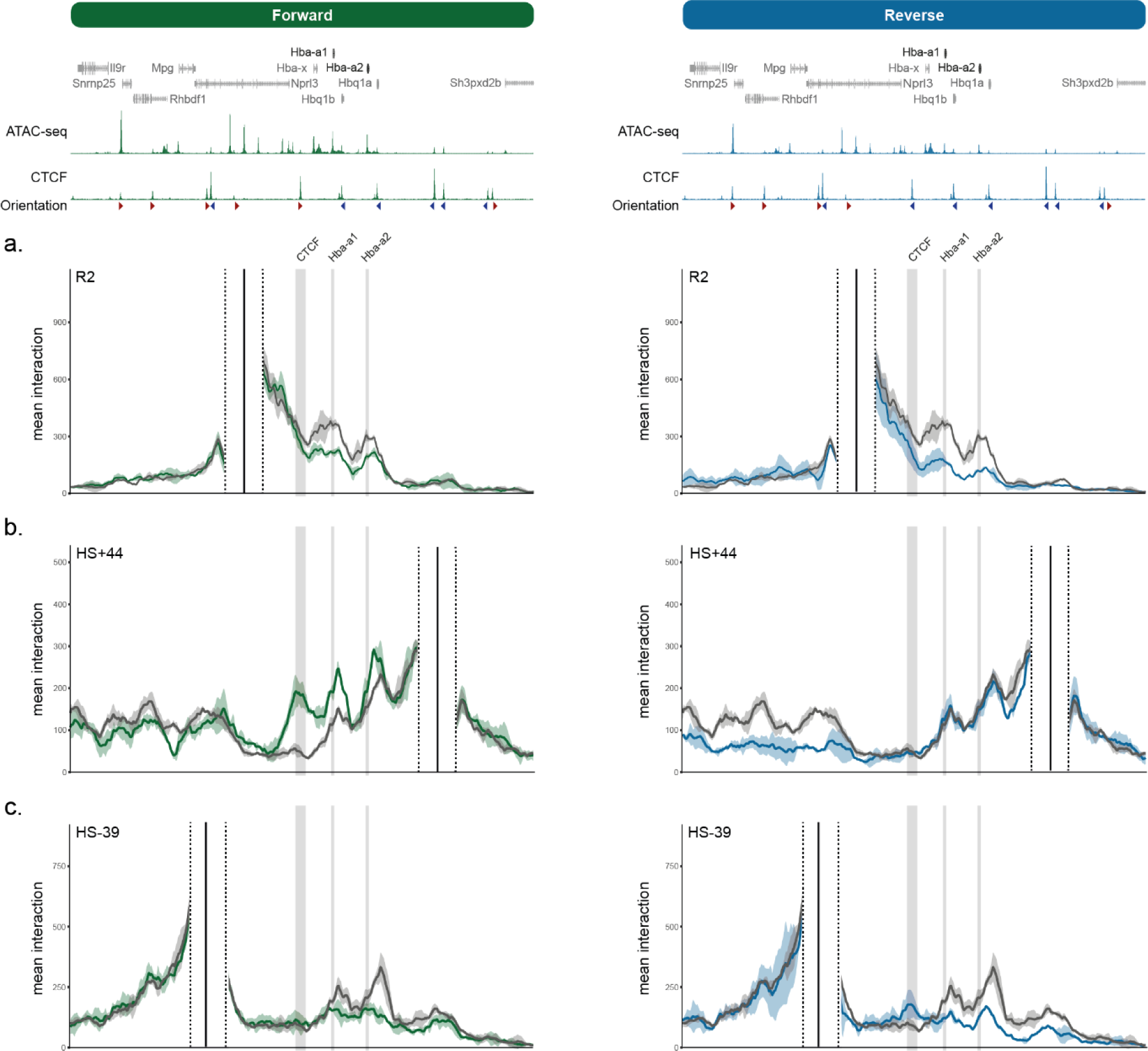
Changes in three-dimensional genome organisation upon introduction of an ectopic CTCF binding site. Top tracks show ATAC-seq and ChIP-seq in the Forward and Reverse EB models. Capture profiles from the viewpoint of R2 (a), CTCF binding sites HS+44 (b) and HS- 38 (c). The profiles show normalised and averaged interactions with a halo representing one standard deviation from biological replicates, for WT (grey, n=3), Forward (green, n=3) and Reverse (blue, n=2) EB-derived erythroid cells. The position of the alpha-globin genes and the ectopic CTCF site is marked in grey. The exclusion zones around capture probes are marked by sashed black lines and the position of the viewpoint by a sold black line.

We next asked if the ectopic insertion altered the looped structure of the flanking CTCF sites, by viewing interactions from the flanking regions. In wild-type cells, the downstream CTCF binding site HS+44 is found in close proximity to nearby CTCF binding sites at the promoters of the theta-globin pseudogenes (*Hbq1a* and *Hbq1b*) and to the upstream boundary elements (HS-38, HS-39, HS-59 and HS-71; Figure 6b). When inserted in the forward orientation, the ectopic CTCF element is convergent with the downstream CTCF sites and is found in close proximity to them: when capturing interactions from HS+44 we observed a new peak of interaction with the ectopic site (Figure 6b). In addition, the interactions with CTCF sites in between the ectopic site and HS+44 increase in the Forward model. In the Reverse model, there is no new peak with the ectopic CTCF binding site, nor is there an increase in other interactions. However, interactions across the locus with upstream CTCF binding sites are reduced (Figure 6b).

Next, we examined interactions from one of these upstream binding sites, HS-39, which marks the 5’ boundary of the alpha-globin sub-TAD. In the Reverse model, there is a modest interaction peak between HS-39 and the ectopic CTCF binding site (Figure 6c). At the same time, interactions between HS-39 site and CTCF sites downstream of the new site are reduced. In the Forward model, interactions with CTCF sites around the alpha-globin genes are reduced, potentially because these are now more frequently interacting with downstream binding sites, but otherwise most interactions remain intact (Figure 6c).

In summary, the ectopic CTCF site is found in close proximity in the Forward and Reverse model with flanking CTCF binding sites that are in convergent orientation, consistent with many previous observations(Rao, Huntley et al. 2014, De Wit, Vos et al. 2015, Guo, Xu et al. 2015). It reduces enhancer-promoter interactions in both models, and is associated with changes in the three-dimensional genome organisation in an orientation-dependent manner. Overall, these experiments support loop-extrusion as a molecular mechanism underlying enhancer-promoter interactions and cohesin as the effector, translocating primarily from the enhancers, and intercepted by boundaries, in a position- and orientation-dependent manner.

## Discussion

Since long-range regulation was first reported, there have been many attempts to explain how enhancers communicate with promoters lying 10s-1000s kb away in the genome(Furlong and Levine 2018). The first models included: tracking, in which a protein such as RNA Polymerase II might be loaded at enhancers and translocate along chromatin to reach and activate their associated promoters(Blackwood and Kadonaga 1998); linking, in which chromatin associated factors arranged along the chromatin fibre might connect the enhancer and promoter(Engel and Tanimoto 2000, Deng, Lee et al. 2012); relocation, in which enhancers and promoters migrate to pre-formed transcription factories(Jackson, Iborra et al. 1998); and passive looping in which linked enhancers and promoters meet frequently by diffusion because they are tethered to each other. Once in contact, by whatever means, the enhancer and promoter might be found in a sub-nuclear structure with a high concentration of transcription factors and co- factors creating a favourable environment for transcription (as reviewed in(Furlong and Levine 2018, Grosveld, van Staalduinen et al. 2021)).

All of these models have been tested experimentally, refined and elaborated upon. Most importantly, in the past decade it has been proposed that regulatory elements within TADs and sub-TADs may be brought into close proximity by the process of cohesin-mediated chromatin loop extrusion(Sanborn, Rao et al. 2015, Fudenberg, Imakaev et al. 2016). This process is delimited by specifically orientated CTCF binding sites that demarcate the active domain(Rao, Huntley et al. 2014, Guo, Xu et al. 2015). Although this is a very compelling and elegant model, experiments in which cohesin has been acutely degraded have been reported to show remarkably few changes in gene expression(Nora, Goloborodko et al. 2017, Rao, Huang et al. 2017, Schwarzer, Abdennur et al. 2017, Wutz, Várnai et al. 2017) and consequently there remains an unresolved discussion about the role of loop extrusion in gene regulation.

Here, we have investigated the role of cohesin and loop extrusion in the enhancer-mediated gene activation at the well-studied mouse alpha-globin locus. We found that the expression of alpha-globin is sensitive to cohesin degradation specifically at earlier stages of differentiation. We further explored the role of translocating cohesin by introducing an ectopic CTCF binding site between the alpha-globin enhancers and genes, which blocks translocating cohesin throughout erythroid differentiation. By introducing the binding site in two orientations, either posed to block cohesin coming from the enhancers or the promoters, we observed lower levels of alpha-globin expression in both models, but a stronger reduction when it predominantly blocks cohesin translocating from the enhancers. This coincides with higher levels of the cohesin subunit Rad21 at the ectopic site and a larger reduction in enhancer-promoter interactions.

Our observations that cohesin depletion does not affect alpha-globin expression late in erythroid differentiation resembles similar experiments in which few changes in gene expression were observed(Rao, Huang et al. 2017, Wutz, Várnai et al. 2017). However, there are potentially confounding factors in the previous experiments that have not been fully explored and which have been further investigated here. First, most previous experiments have been conducted in steady-state conditions(Symmons, Uslu et al. 2014, Fudenberg, Abdennur et al. 2017, Luppino, Park et al. 2020, de Wit and Nora 2023). However, of interest, in experiments analysing the effect of cohesin depletion in primary macrophages in response to inflammatory stimuli there was a large effect on inducible genes whereas expression of most constitutive genes was unchanged(Cuartero, Weiss et al. 2018, Robles-Rebollo, Cuartero et al. 2022). During erythropoiesis many genes are induced via enhancer-mediated activation and the globin genes provide a clear example of this. We have shown here that, when cohesin is depleted, globin RNA is downregulated in earlier erythroid cells specifically (day 5 and 6 EB-derived erythroid cells) whereas the widely expressed genes immediately surrounding them are not affected (Supplementary figure 2). It is possible that the lack of effects of cohesin depletion on gene expression seen in previous experiments reflects the fact that loop extrusion is particularly important for establishing enhancer–promoter communication during differentiation or in response to stimuli, and that when established, other factors, such as protein-protein contacts, maintain the interaction in a steady state. Alternatively, after activation, transcription may be maintained by re-initiation rather than requiring enhancer-driven activation.

Another consideration is the degree to which cohesin is downregulated in these experiments. In some experiments, in contrast to the global loss of TADs and CTCF loops, in the absence of cohesin(Rao, Huang et al. 2017, Schwarzer, Abdennur et al. 2017, Wutz, Várnai et al. 2017), enhancer–promoter interactions are not completely abolished(Thiecke, Wutz et al. 2020, Liu, Maresca et al. 2021, Aljahani, Hua et al. 2022, Calderon, Weiss et al. 2022). In line with these observations, it is of interest that two recent reports based on analysis with Micro-C(Hsieh, Cattoglio et al. 2022) and a targeted MNase-based 3C(Goel, Huseyin et al. 2023) did not detect clear changes in enhancer–promoter interactions upon cohesin depletion in mES cells. It is possible that these different reports reflect the different degrees to which cohesin may have been depleted, which has previously been observed with similar chromatin-bound proteins such as CTCF(Nora, Goloborodko et al. 2017, Khoury, Achinger-Kawecka et al. 2020).

We have also used an orthogonal approach to depleting cohesin, by placing an ectopic CTCF binding site between the enhancers and target genes to block loop extrusion. We found that a single CTCF binding site was sufficient to disrupt enhancer function considerably. This is in contrast to some other studies, were multiple CTCF sites had to be inserted in tandem to reach similar levels of insulation(Huang, Zhu et al. 2021). This partly reflects that HS-38 is a strong insulator, however, binding site sequence alone cannot adequately explain this. Several of the same CTCF binding site sequences that did not perform as strong insulators when inserted at the Sox2 locus(Huang, Zhu et al. 2021), did act as strong boundaries in the assay at the alpha-globin locus(Tsang, Stolper et al. 2023). We therefore speculate that locus-specific characteristics, such as the dependence of a gene on its enhancer for expression or the distance between the enhancer and promoter, are important variables when evaluating the effect of a boundary element. Others have suggested that the role of loop extrusion is limited to very long-range enhancer promoter interactions(Calderon, Weiss et al. 2022, Rinzema, Sofiadis et al. 2022), but the observations made here of the effect of single or multiple CTCF sites insertions as well as the cohesin depletion data, indicates a role for loop extrusion at the relatively short (30-50 kb) chromosomal interaction between the alpha-globin enhancers and promoters.

We also observed a striking difference between models with a forward or reverse oriented ectopic CTCF binding site. The difference is most easily explained by a mechanism that involves tracking along the chromatin fibre and is therefore in accord with loop extrusion. Any mechanism not involving tracking along the chromatin fibre would be agnostic to the binding site orientation. Fundamentally, the Forward and Reverse models test the effect of blocking loop extrusion from different directions. Viewed in this light, the unequal decrease in alpha-globin expression between the Forward and Reverse CTCF sites indicates that loop extrusion coming from enhancers plays a more important role in setting up enhancer-promoter interactions. It is possible that cohesin originating from the enhancers somehow plays a role in enhancer function, and therefore blocking extrusion from this direction has an especially strong effect on gene expression. It is also possible that cohesin extrusion is not equal in both directions along the chromatin fibre.

The data presented here are consistent with previous work suggesting that active enhancers may provide a loading site for cohesin which then translocates and thereby juxtaposes the enhancers to flanking promoters. Previous evidence of enhancers as sites of cohesin loading was based on the observations that more cohesin is present when an enhancer cluster is active(Hua, Badat et al. 2021), or on the fact that cohesin levels decrease when a super enhancer is deleted(Vos, Valdes-Quezada et al. 2021). However, these models cannot distinguish between cohesin loading at active loci, which include enhancers, promoters and gene bodies, and loading at enhancers specifically. Indeed, NIPBL, which is necessary for the loading of cohesin onto the chromatin fibre, is present at both active enhancers and active genes(Hua, Badat et al. 2021), although there is some evidence that its presence at active genes could be a technical artefact(Banigan, Tang et al. 2023). There have also been reports of interactions between NIPBL and mediator(Kagey, Newman et al. 2010, Mattingly, Seidel et al. 2022), which is present at the alpha-globin enhancers. Cohesin predominantly translocating from the enhancers also fits with the observations of higher levels of Rad21 at the reverse oriented binding site.

From current observations it seems that the probability of an active distal enhancer interacting with the surrounding promoters will depend on many dynamic factors which will differ for different loci(Furlong and Levine 2018). These currently include: location within a shared TAD and particularly within a shared sub-TAD(Symmons, Uslu et al. 2014); the distance between the enhancer and promoter(Zuin, Roth et al. 2022); the “strength” of the enhancer(Long, Prescott et al. 2016, Catarino and Stark 2018); the “compatibility” between enhancer and promoter(Zabidi, Arnold et al. 2015, Martinez-Ara, Comoglio et al. 2022); competition between surrounding promoters(Cho, Xu et al. 2018); and the homotypic and heterotypic interactions between proteins which contribute to the interacting elements and the transcriptional hub(Deng, Lee et al. 2012). The data presented here and elsewhere, are consistent with a model in which the degree of cohesin-mediated loop extrusion and the direction of translocation also play a significant role in establishing enhancer-promoter interactions especially during differentiation or in response to stimuli rather than in the steady state(Cuartero, Weiss et al. 2018).

We conclude that the data presented here are consistent with a model in which the alpha-globin enhancers and promoters are brought into proximity by a tracking mechanism involving cohesin-mediated loop extrusion, and will contribute to the hypothesis that enhancers specifically play a strong role in directing loop extrusion and gene expression.

## Methods

### Cells and genome editing

E14Tg2a mouse embryonic stem (ES) cells were used for all mouse ES cell experiments in this study. The cells were maintained in KnockOut™ DMEM or Glasgow’s Minimal Essential Medium supplemented with 10% fetal bovine serum, 1 mM sodium pyruvate, 2 mM L-Glutamine, 1x non-essential amino acids (all Gibco), beta-mercaptoethanol, and leukemia inhibiting factors. During gene targeting, the media was supplemented with penicillin and streptomycin (Gibco). Cells were usually grown on 0.1% (v/v) gelatine (Sigma-Aldrich: G1393), except during gene targeting for the CTCF insertion mouse model and prior to blastocyst injection, when they were grown on mitotically inactivated mouse embryonic fibroblasts (MEFs).

For the CTCF insertion cells (Forward/Reverse model) and mouse model, a double nickase CRISPR approach was used. Single guide-RNA plasmids were generated by cloning oligonucleotides to the target site (see table 1) into pX335 (Addgene plasmid no. 42335(Cong, Ran et al. 2013)). pX335 had been modified to contain a puromycin selection cassette. The homology direct repair (HDR) plasmid was prepared by amplifying 1.4 and 1.2 kb homology arms from genomic DNA, which were then cloned into the pJET vector (CloneJET PCR cloning kit, Thermo Scientific). A synthetic DNA fragment (Genewiz) containing the 82 bp CTCF binding site, flanked by heterospecific lox sites (loxP and lox2272(Lee and Saito 1998)), two FRT sites and flanking rox sites was cloned between the homology arms (see supplementary table 1). A neomycin selection cassette was cloned between the FRT sites and the sequence of the final plasmid sequence was confirmed by Sanger sequencing.

**Table 1:**
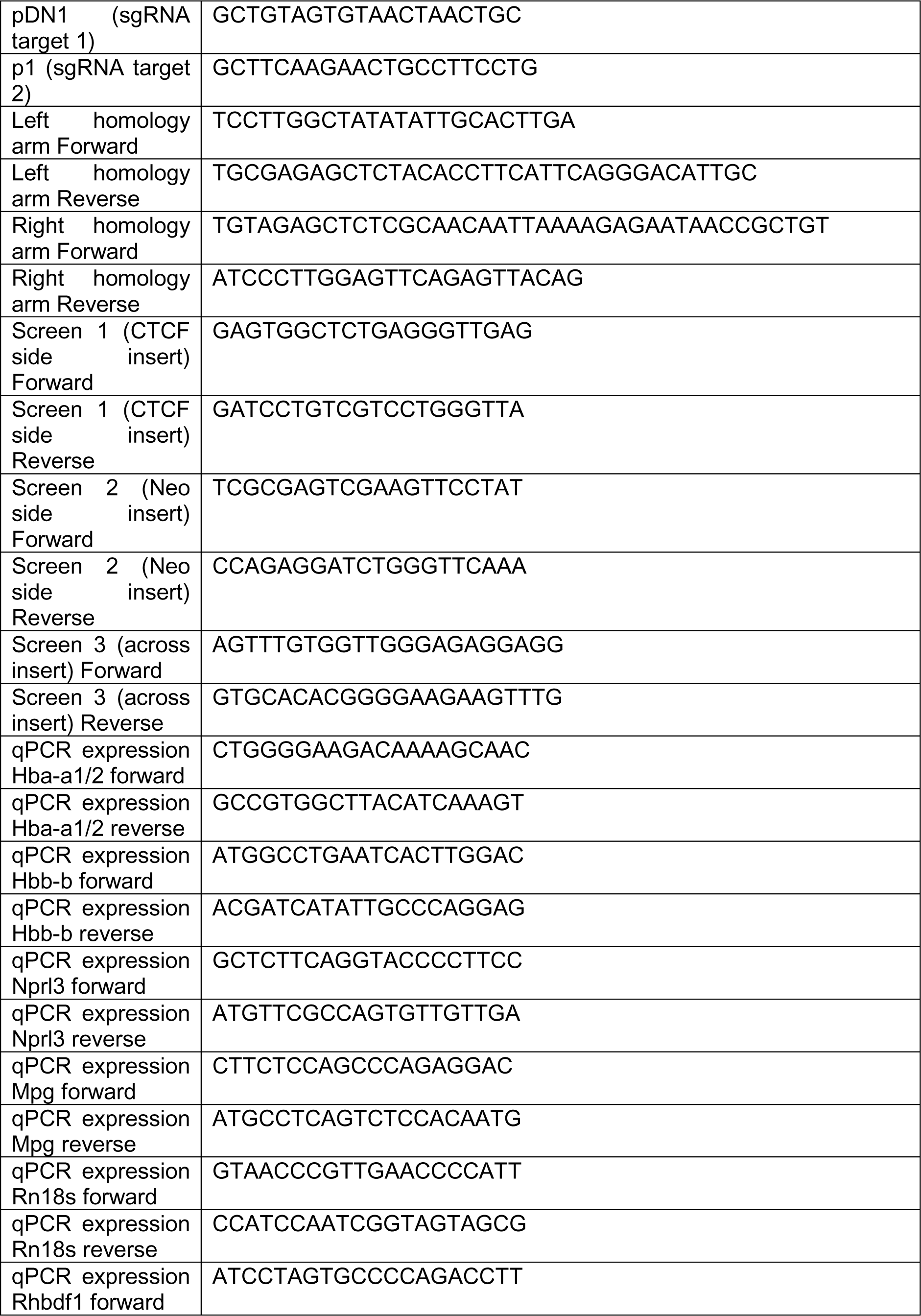
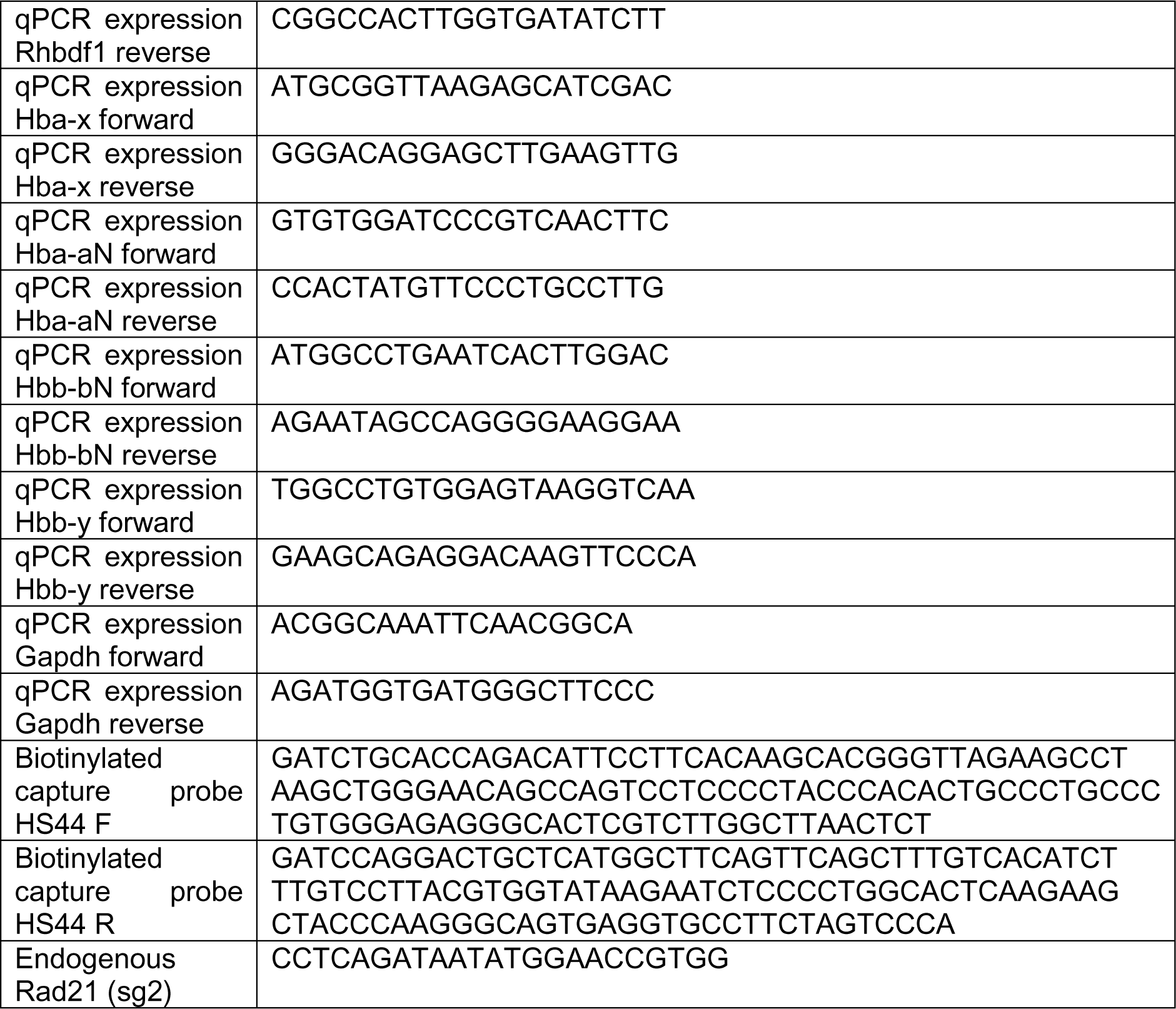
DNA sequences used in this study.

Wild-type mouse ES cells were electroporated using the Neon transfection system (Invitrogen) using 3 × 1400V for 10 ms with 2.5 μg of the two sgRNA nickase-Cas9 plasmids and 5 μg targeting vector. The transfected ES cells were plated at low density and selected with 600 ng/ml puromycin for two days, followed by 350 ng/ml G418 selection for 5 days. Individual clones were selected, expanded and initially screened for correct integration using homology-arm spanning PCRs (Immolase DNA polymerase, Bioline, supplemented with Q solution, Qiagen). Genotypes of positive clones were confirmed by Sanger sequencing. Three clones with the correct insertion were transiently transfected with either 5 μg pFlpO (Addgene #13793) to remove the neomycin selection cassette or both 5 μg pFlpO (Addgene #13792) and 5 µg of pCrePac(Taniguchi, Sanbo et al. 1998) to simultaneously remove the selection cassette and invert the CTCF binding site. This provided clones with the forward and reverse orientated CTCF site in triplicate.

For the CTCF activity reporter system, genetically modified mESC models were generated using the double nickase HDR strategy. In brief, the two single guide-RNAs targeting the insertion site were individually cloned into the pX335 vector containing puromycin selection cassette as described above. To generate the HDR donor plasmid containing the inserted CTCF elements, oligonucleotides corresponding to each CTCF element were cloned into the pROSA-TV2 vector (created by Prof. Benjamin Davies group) between the pair of BsaI site with the NEB Golden Gate Assembly Kit (NEB #E1601). The pROSA-TV2 was designed to contain the 1.4 and 1.2 homology arms of the inserted site and a hygromycin selection cassette that is flanked by a pair of lox sites (loxP). Sequences of the CTCF element oligonucleotides are listed in the supplementary table 1. The mESC E14TG2a used for the CTCF activity reporter assays was modified by hemizygously deleted one wild-type copy of the alpha-globin locus to facilitate the further genetical modifications. In addition, a yellow fluorescent protein (YFP) was tagged to the *Hba-a1* gene, so that the YFP level is used as a readout for the Hba-a1 expression level, as reported previously(Francis, Harold et al. 2022). The mESCs were co-transfected with the 1.66 μg of the two sgRNA nickase-Cas9 plasmids and 0.83 μg of the HDR plasmids using the Lipofectamine LTX and Plus Reagent (Invitrogen #15338-100). Transfected cells were first selected by 1 μg/ml puromycin for 2 days and subsequently selected by 250 μg/ml hygromycin for 6 days. The drug selected cells were then transfected with the Cre-recombinase vector to remove of the integrated hygromycin-resistance cassette. Genetically modified mESCs were FACS sorted into 96-well plate and grown into individual colonies. The colonies were screened for the correct integration by homology-arm spanning PCRs and the genotypes were further confirmed by Sanger sequencing.

For the Rad21 dTAG experiments, the native Rad21 gene was tagged with FKBP12^F36V^ (Nabet, Roberts et al. 2018) by CRISPR-Cas9 mediated HDR in wild-type mouse ES cells. To achieve this, the sgRNA (see table 1) was cloned in to pX458 (Addgene plasmid no 48138(Ran, Hsu et al. 2013)), which encodes Cas9 and enhanced green fluorescent protein (EGFP). HDR templates containing the FKPB12^F36V^ tag were synthesised (see supplementary table 1) and cloned by GeneArt (Invitrogen) into double-stranded plasmid vectors (pMX).

Wild-type mouse ES cells were transfected using Lipofectamine LTX Reagent with PLUS Reagent (Invitrogen), which were performed according to manufacturer’s instructions: 5 μl LTX reagent, a total of 2.5 μg purified plasmid DNA (1:3 ratio of sgRNA/Cas9 plasmid DNA:donor plasmid DNA), 2 μl PLUS reagent, and 250 μl Opti-MEM (Gibco). After 24 hours, transfected cells were rinsed with PBS and given fresh media. Transfected cells were selected based on the presence of EGFP by FACS after 48 hours and positive cells were single cell sorted into gelatine coated 96 well tissue culture plates. Following 10 days of growth, surviving colonies were spit across replica plates and screened for positive clones by PCR.

### Animal procedure

The mutant mouse strains reported in this study were backcrossed and maintained on a C57BL/6J background. Protocols were approved through the Oxford University Local Ethical Review process. Experimental procedures were in accordance with the European Union Directive 2010/63/EU and/or the UK Animals (Scientific Procedures) Act (1986), and reviewed by the clinical medicine Animal Welfare and Ethical Review Body (AWERB). Experimental procedures were conducted under project licences PPL 30/2966 and PPL 30/3339.

ES cells with the original insertion (as described above), prior to Flp and Cre recombinase treatment, were used for blastocyst injection. After germline transmission, these were crossed with a Flp-expressing mouse line (B6.Cg-Tg(ACTFLPe)9205Dym/J; Jackson Laboratory stock #005703), and following positive genotyping, with a Cre-expressing line(Mao, Fujiwara et al. 1999) to produce both a forward and reverse CTCF insertion strains.

### Fetal liver culture

The *in vitro* mouse fetal liver culturing system is based on previous protocol (von Lindern, Deiner et al. 2001, Chen, Liu et al. 2009). Briefly, E12.5 mouse fetal liver were dissected and cultured in StemPro medium (Invitrogen) supplemented with Epo (1 U/mL; Janssen, PL 00242/029), SCF (50 ng/mL; Peprotech, 250-03), dexamethasone (1 μM; Hameln, DEXA3.3) and 1× L-Glutamine (Invitrogen) for 6–8 days to expand the erythroid progenitor population. During culture the erythroblast cell count was maintained at 1 × 106 cells/ml or under. If necessary cells were frozen at day 4 and after thawing expanded for a further 2-4 days.

### In vitro embryoid body formation and differentiation

An *in vitro* haematopoietic differentiation protocol(Francis, Harold et al. 2022) was used to differentiate mouse ES cells into embryoid bodies (EBs), based on a previously published protocol(Keller 1995). In brief, proliferating ES cells were cultured in IMDM-ES media (Iscove’s Modified Dulbecco Medium (IMDM) (Gibco) supplemented with 15% fetal bovine serum (FBS), 100 U/ml penicillin/streptomycin, 1000 U/ml LIF, and 1.5 × 10^-4^ M monothioglycerol (MTG; Sigma-Aldrich: M6145)) for 20 to 48 hours prior to primary plating. Subsequently, cells were plated in 10 ml differentiation media on bacterial petri dishes (15000-50000 cells/plate) and cultured for 7 days. Differentiation medium consisted of IMDM media supplemented with 15% heat-inactivated FBS (1 hour at 56 °C), 2 mM L-Glutamine, 300 mg/ml transferrin (Roche), 50 g/ml ascorbic acid, 3 × 10^-4^ M MTG and 5% Protein-Free Hybridoma Media II (Gibco).

### Isolation of erythroid cells

After 5, 6 or 7 days of differentiation, EBs were disaggregated using 0.25% trypsin (Gibco) and stained with 0.5 μg FITC Rat anti-mouse CD71 antibody (BD Pharmingen or eBioscience) per 10^6^ cells for 20 minutes at 4 °C, followed by washing and incubation with anti-FITC MACS microbeads (Miltenyi Biotec) for 15 minutes at 4 °C. Stained cells were then purified using MACS lineage selection columns (Miltenyi Biotec) and immediately processed for various assays or treated with dTAG. For dTAG treatment, CD71+ cells were plated in differentiation media, as described above, supplemented with 1U/ml Epo (Janssen, PL 00242/029), and 500 nM dTAG-13 for the treated cells. Control cells were incubated for the same amount of time as the treated cells. The purity of the isolated erythroid population was verified by fluorescent-activated cell sorting (FACS; Attune Nxt Flow Cytometer (Thermo Fisher)).

After 6 to 8 days of expansion, fetal liver cells were stained with 1 μg anti-mouse Ter119-PE antibody (BD Pharmingen) per 10^6^ cells for 20 minutes at 4 °C. Cells were washed with 10% FBS in PBS and resuspended in 135 ml cold PBE buffer (PBS with 0.5%BSA and 2 mM EDTA) with 15 ml Ter119-PE microbeads (Miltenyi Biotec) per 10^7^ cells, and incubated for 15 minutes at 4 °C. Erythroid cells were then isolated using MACS lineage selection columns (Miltenyi Biotec) and processed for various assays. Purity of isolated populations was assessed as described above.

Primary erythroid cells were obtained from the spleen of adult mice treated with phenylhydrazine, as described previously(Spivak, Toretti et al. 1973). Spleens were mechanically disrupted in 10% FBS in PBS to produce a single cell suspension. Cells were resuspended in 1 ml 10% FBS in PBS and stained with 6 μg anti-mouse Ter119-PE antibody (Miltenyi Biotec) per 10^8^ cells for 20 minutes at 4 °C. After washing with 10% FBS in PBS, cells were resuspended in 8 ml cold PBE buffer (PBS with 0.5% BSA and 2 mM EDTA) and 2 ml anti-PE microbeads (Miltenyi Biotec) per 10^8^ cells and incubated for 15 minutes at 4 °C. Ter119+ cells were selected with MACS lineage specification columns (Miltenyi Biotec), typically using one column for up to 5 × 10^8^ cells, and processed for various assays Purity of isolated populations was assessed as described above.

### Western blot

Total protein from 500,000 EB-derived Cd71+ cells were harvested for western blotting. Cells were washed in ice-cold PBS twice and lysed in RIPA buffer (Sigma #SLCG2240) supplemented with 10% protease inhibitor cocktail (Roche, 11873580001) for 30 minutes at 4°C. Protein lysates were quantified using Qubit Protein Assay Kit (Invitrogen #Q33211) and mixed with the western blot denaturing loading dye (125mM Tris-HCl, 4% SDS, 50% Glycerol, and 0.4g Orange G) at a 1:10 ratio. DTT (Sigma #43816) was added at a final concentration of 100 mM and the samples were denatured at 95°C for 10 minutes. 40 μg of protein were loaded and run on pre-casted 4-12% NuPAGE Bis-Tris gels (Invitrogen #NP0335) for 90 minutes at 120V. Samples were transferred on nitrocellulose membrane (Amersham GE Healthcare #10600008) for 90 minutes at 30V. The membrane was blocked in 5% milk in PBS for 1 hour at room temperature with constant agitation. The membrane was incubated with primary antibodies at 4°C overnight. The dilution for the mouse anti-Flag antibody (Sigma, F1804) is 1:1000 and the dilution for the rabbit anti-Beta actin antibody (Abcam #ab115777) is 1:200 in blocking buffer with 0.2% Tween-20. The membrane was washed three times with PBS with 0.1% Tween-20 and was incubated with secondary antibodies for 1 hour in dark. The dilutions of the near-infrared coupled secondary antibodies IRDye 800CW goat anti-rabbit IgG (Abcam, ab216773) and IRDye 680RD goat anti-mouse IgG (Abcam #ab216776) are 1:1000 in blocking buffer with 0.2% Tween-20. The membrane was washed three times with PBS-0.1% Tween-20 before detection with the iBright FL1000 scanner and software (Invitrogen) on the 680nm and 800nm channel. The quantification of the protein bands on the western blot was performed by using the Image J package and the result reflects the amount of the Flag-Rad21 protein relative to the Beta-actin loading control as a ratio.

### Expression analysis

Total RNA was extracted from 5×10^5^ – 5×10^6^ cells. Cells were lysed in TRI Reagent (Sigma-Aldrich) according to manufacturer’s instruction and immediately stored at -80 °C. Total RNA was isolated either via chloroform extraction or using a Direct-zol MiniPrep or MicroPrep kit (Zymo Research).

For extraction using chloroform, chloroform was added to the sample in TRI reagent in a ratio 1:5 (v/v) and the sample was mixed vigorously, followed by room temperature incubation. Then the sample was centrifuged (15000 RPM, 4 °C) for 10 minutes and the aqueous layer was transferred and precipitated with isopropanol and glycoblue (Thermo Fisher) at room temperature for a few minutes. The samples were centrifuged (15000 RPM, 4 °C, 10 minutes) and the pellet containing the RNA was washed with 70% ethanol, air dried and resuspended in DNAse/RNase-free water. Samples were then treated with DNase using the TURBO DNA-free kit (Invitrogen) according to manufacturer’s instruction. Briefly, samples were treated with DNase enzyme for 30 minutes followed by inactivation for 5 minutes, after which the samples were centrifuged and the supernatant was used for cDNA synthesis after quality control steps. RNA isolated using the Direct-zol kits was performed according to manufacturer’s instructions, but the DNase I treatment was extended to 30 minutes at room temperature (rather than 15 minutes). The quality of all isolated RNA was assessed using RNA Analysis Screentape (Tapestation, Agilent Technologies) and only samples with high (>7.5) RNA integrity number (RIN) scores were used in subsequent steps.

Up to 1 μg RNA was used for cDNA synthesis using the Superscript III Reverse Transcriptase kit (Invitrogen) following manufacturer’s instructions. For each sample, a control without reverse transcriptase enzyme was included to account for any incomplete genomic DNA digestion. cDNA was diluted 5× before qPCR analysis to determine the relative changes in gene expression. qPCR analysis was performed in triplicate using Fast SYBR green master mix (Applied Biosystems) using primers listed in Table 1. The ΔΔCt-method was used for relative quantification of RNA abundance.

### ATAC-seq

Assay for transposase-accessible chromatin (ATAC)-seq was performed as previously described(Buenrostro, Giresi et al. 2013). Briefly, 60 000 – 75 000 cells were lysed in cold lysis buffer (10 mM Tris HCl, 10 mM NaCl, 3.4 mM MgCl_2_, 0.1% IGEPAL CA-630) to extract nuclei. Pelleted nuclei were incubated in transposase reaction mix (1 X TD reaction buffer, 2.5 µl Tn5 transposase (Nextera, Illumina)) for 30 min at 37 °C. Tagmented DNA was purified using a MinElute PCR purification kit (Qiagen) and amplified and indexed with NEBNext High-Fidelity 2X PCR Master Mix (NEB) and custom primers. Amplified libraries were purified with PCR cleanup kit (Qiagen) and library quality was assessed on a Tapestation (Agilent Technologies). Libraries were sequenced on the Illumina NextSeq platform using a 75-cycle paired end kit (NextSeq 500/550 High Output Kit).

### ChIP-seq

A Chromatin Immunoprecipitation (ChIP) Assay Kit (Merck Millipore 17-295) was used for all CTCF ChIP experiments following manufacturer’s instructions. First, chromatin of 5-10×10^6^ isolated erythroid cells was fixed in 1% formaldehyde for 10 min. The crosslinking reaction was quenched with 125 mM glycine for 5 min. Chromatin was sonicated using the Bioruptor or Bioruptor pico (Diagenode) to produce fragments of 200-500 bp. Sonicated chromatin was incubated with 10 µl anti-CTCF antibody per sample (Millipore 07-729) at 4 °C on a rotator overnight. Protein A agarose beads were added and incubated with the chromatin for 60 minutes at 4 °C with rotation, and beads were sequentially washed and eluted following manufacturer’s instructions.

For Rad21 ChIP, samples were either performed using the Millipore kit as described previously(Hanssen, Kassouf et al. 2017) or using a magnetic-bead based protocol. Briefly, 10^7^ cells were fixed with 2 mM disuccinimidyl glutarate (DSG; Thermo Fisher) for 50 minutes, followed by 1% formaldehyde for 10 minutes. The crosslinking reaction was quenched with a final concentration of 125 mM glycine, for 5 minutes. After cell lysis, chromatin was sonicated using the Bioruptor (Diagenode), Bioruptor pico (Diagenode) or ME220 Focused-ultrasonicator (Covaris) to produce fragments of 200-500 bp. Sonicated chromatin was incubated with 10 µl Rad21 antibody (Abcam, ab992) per sample at 4 °C on a rotator overnight. Samples were than treated as CTCF ChIPs described above. Alternatively, using a magnetic bead-based protocol, samples were fixed with 1% formaldehyde for 10 minutes at room temperature and quenched with 125 mM glycine for 5 minutes. Cells were lysed in SDS lysis buffer (1% SDS, 10 mM EDTA, 50 mM Tris-HCl pH 8) supplemented with Proteinase Inhibitor Cocktail (Roche, 11697498001) and chromatin was sonicated using the Bioruptor pico (Diagenode) to produce fragments of 200-500 bp. Chromatin was isolated using a Rad21 antibody (ab992) and Protein-G Dynabeads (Thermo Fisher) and washed three times with RIPA buffer (50 mM HEPES-KOH, pH 7.6, 500 mM LiCl, 1 mM EDTA, 1% Igepal CA-630, 0.7% sodium deoxycholate) and two times with Tris-EDTA (TE buffer with 50 mM NaCl). Precipitated chromatin was eluted in 1% SDS, 10 mM EDTA and 50 mM Tris-HCl. After decrosslinking and treatment with RNase A and proteinase K, DNA was purified using DNA Clean-up and Concentration kit (Zymo Research).

Good quality libraries, as determined by qPCR enrichment, were prepared for sequencing with an Ultra II Library Prep Kit (NEB) and sequenced on the Illumina NextSeq platform using a 75-cycle paired end kit (NextSeq 500/550 High Output Kit).

### Capture-C

Capture-C was performed as previously described, either following the next-generation Capture-C protocol(Davies, Telenius et al. 2016) or the NuCapture-C protocol(Downes, Smith et al. 2022). A total of 3 - 10 × 10^6^ EB-derived CD71+ were processed per biological replicate and libraries of sufficient quality were indexed using NEBNext Ultra II DNA Library Prep Kit for Illumina (New England Biolabs: E7645) according to the manufacturer’s instructions. We performed capture enrichment using biotinylated oligos previously published (HS-38 and R2)(Hanssen, Kassouf et al. 2017) or listed below (HS44; table 1). Data was analysed using scripts available at https://github.com/Hughes-Genome-Group/CCseqBasicS. Data visualisation and statistical analysis was performed using CaptureSee(Telenius, Downes et al. 2020).

### ATAC-seq and ChIP-seq data analysis

Next-generation sequencing reads were trimmed using trimgalore (version 0.3.1 https://www.bioinformatics.babraham.ac.uk/projects/trim_galore/) with stringency 2 and aligned with bowtie2(Langmead and Salzberg 2012) (version 2.1.0) to a custom genome based on mm9, edited to include CTCF site insertion as appropriate. Samtools(Li, Handsaker et al. 2009) (version 1.3) was used to remove PCR duplicates and select reads from chromosome 11, and bedgraphs were created using bamCoverage (part of deeptools version 2.2.2(Ramírez, Ryan et al. 2016)), with normalisation set to RPKM. Replicates were merged with unionbedg (bedtools version 2.25.0(Quinlan and Hall 2010)) and bigwig tracks were created with bedGraphToBigWig (deeptools). To compare bigwig files from different genotypes, tracks were shifted to align over genes and other genomic features. Finally, data was visualised using the UCSC genome browser(Kent, Sugnet et al. 2002).

### Read count analysis and peak calling

Reads were counted over the ectopic CTCF site (chr11:32,171,303-32,172,502) for every sample (aligned to CTCFins custom genome). For normalisation, peaks were called on a subset of ChIP samples with Lanceotron(Hentges, Sergeant et al. 2021), using model wide-and-deep_jan-2021, on bigWig files generated using bamCoverage (part of deeptools 3.4.3, (Ramírez et al. 2016)) with extendReads (default), binsize 1 and normalisation set to RPKM, after aligning to mm9. Peaks with a peak score between 0.8 and 1 were used for further analysis. From these peak calls, peaks present in multiple samples were used to create a peak list. Next, bedtools multicov (bedtools v2.29.2) was used to count reads over the peak call list for every sample and the average number of reads per peak was used to normalise read counts over the ectopic CTCF site.

**Supplementary table 1:**
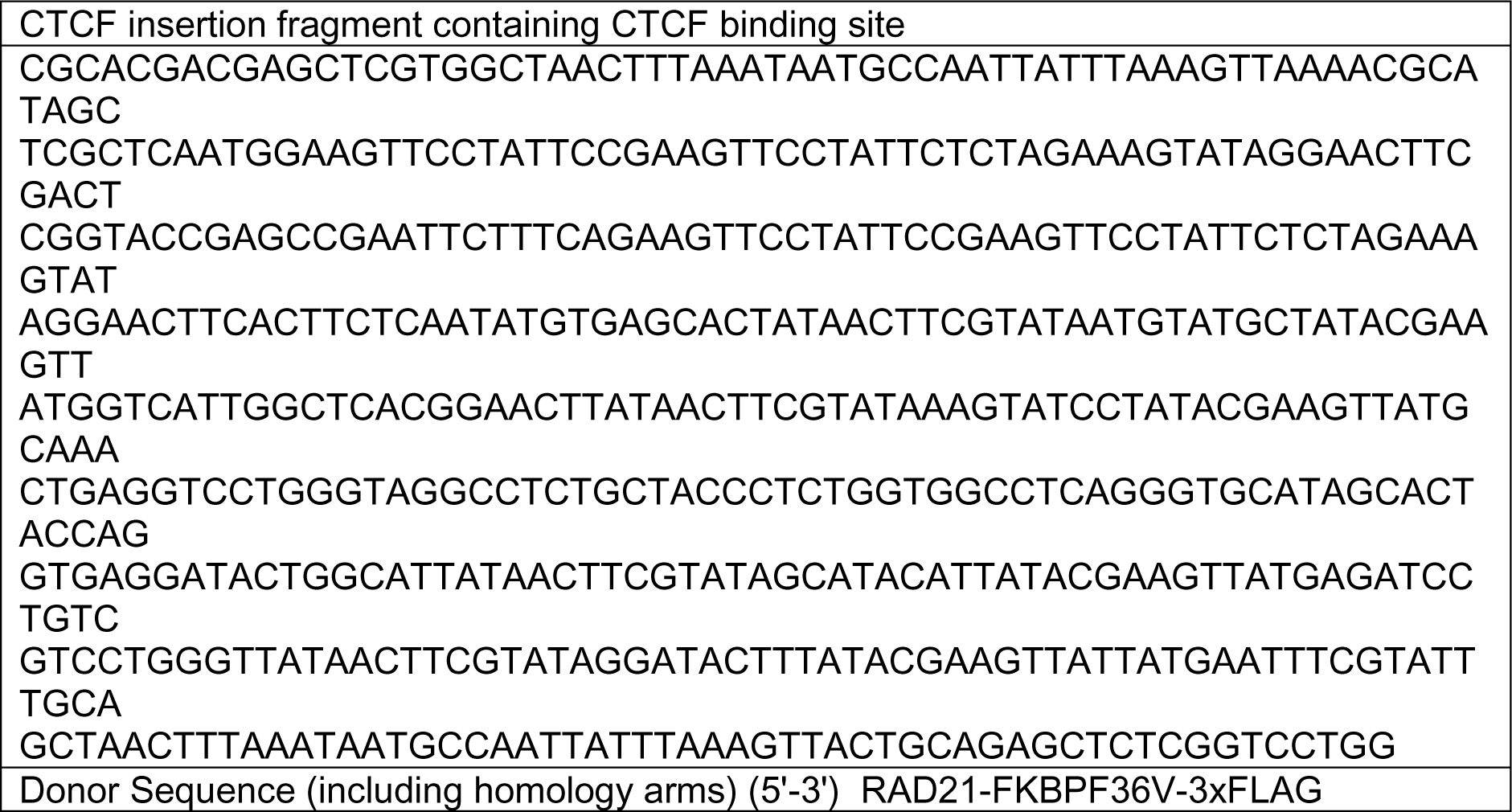
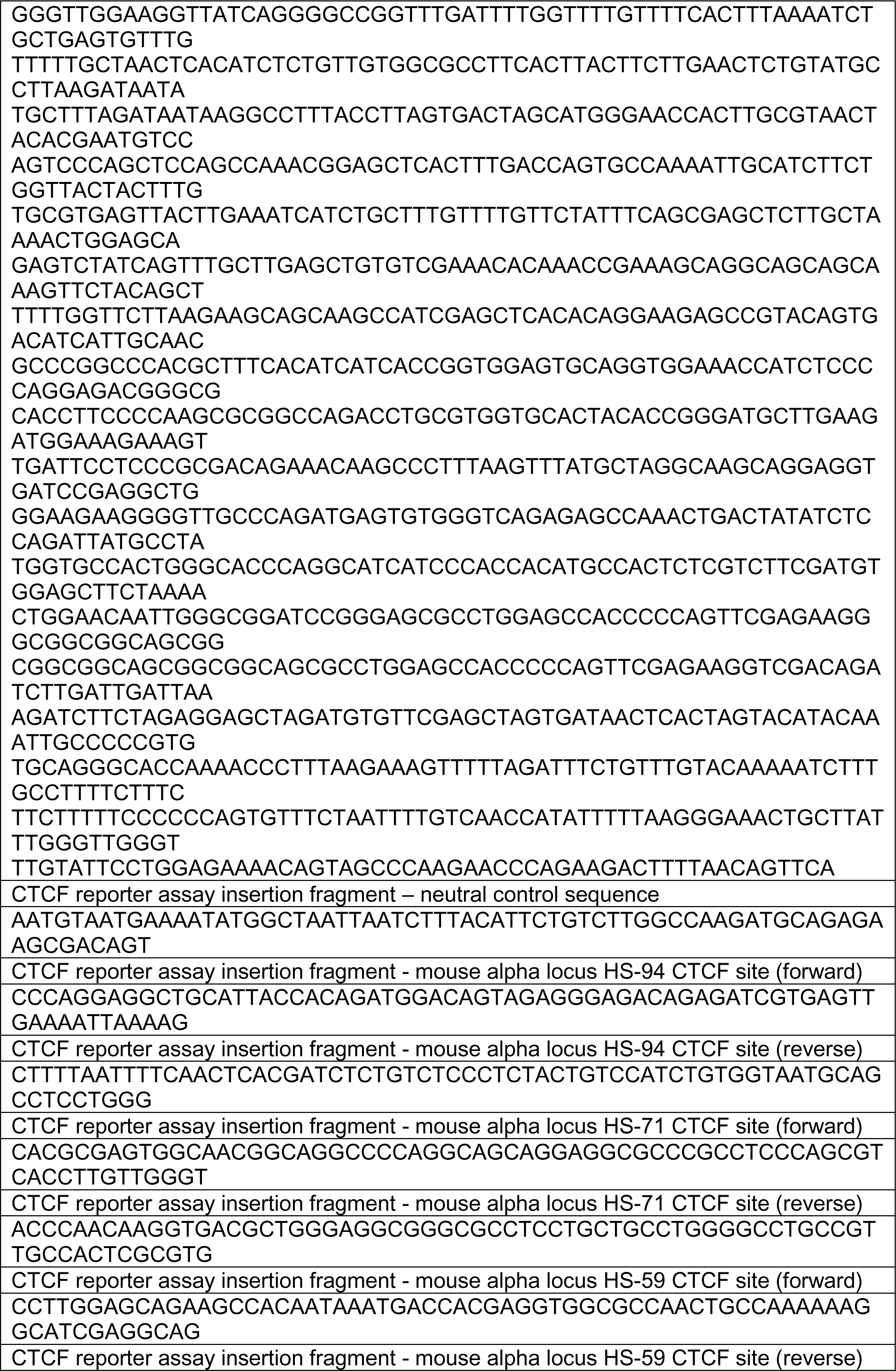
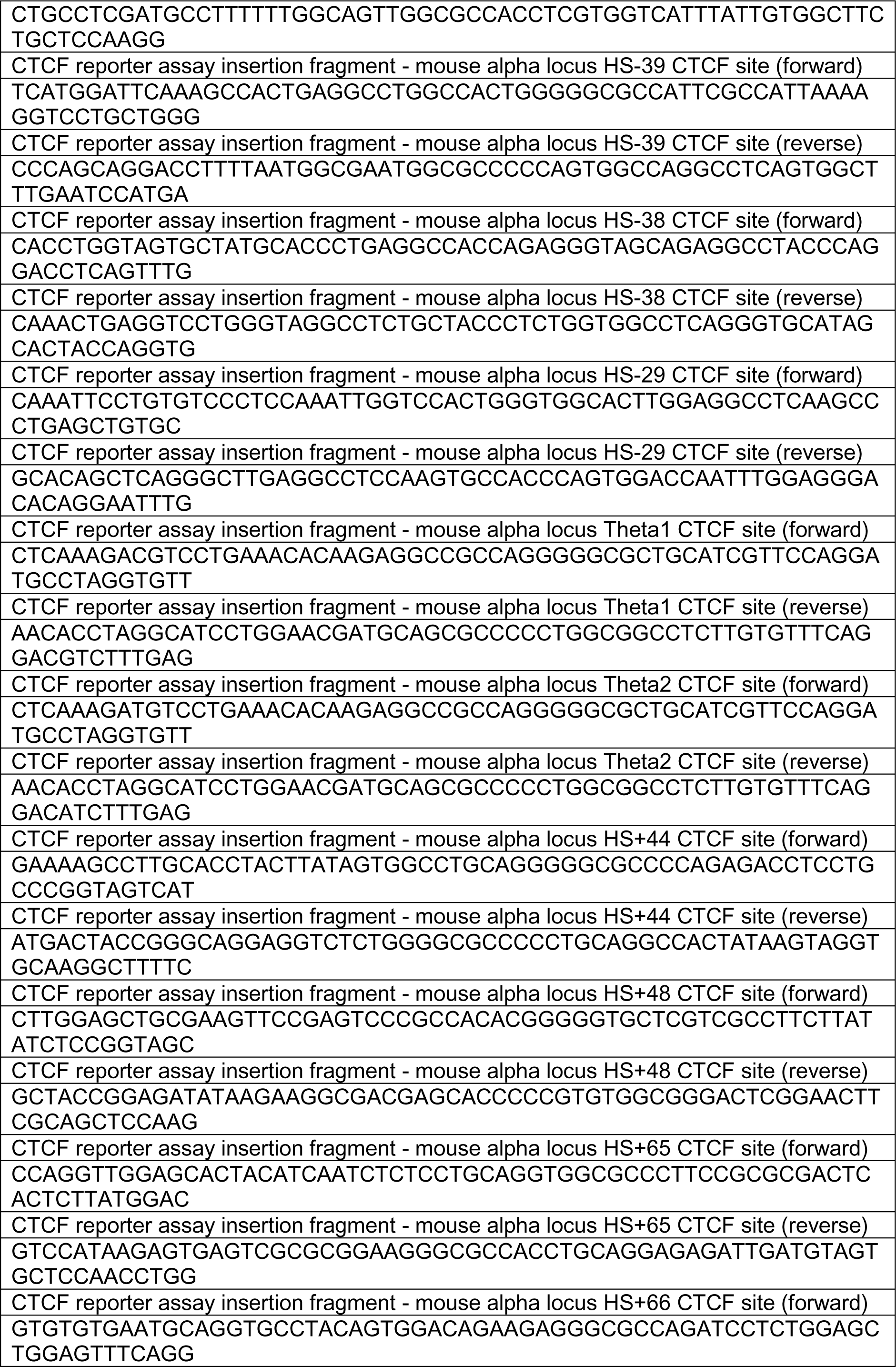
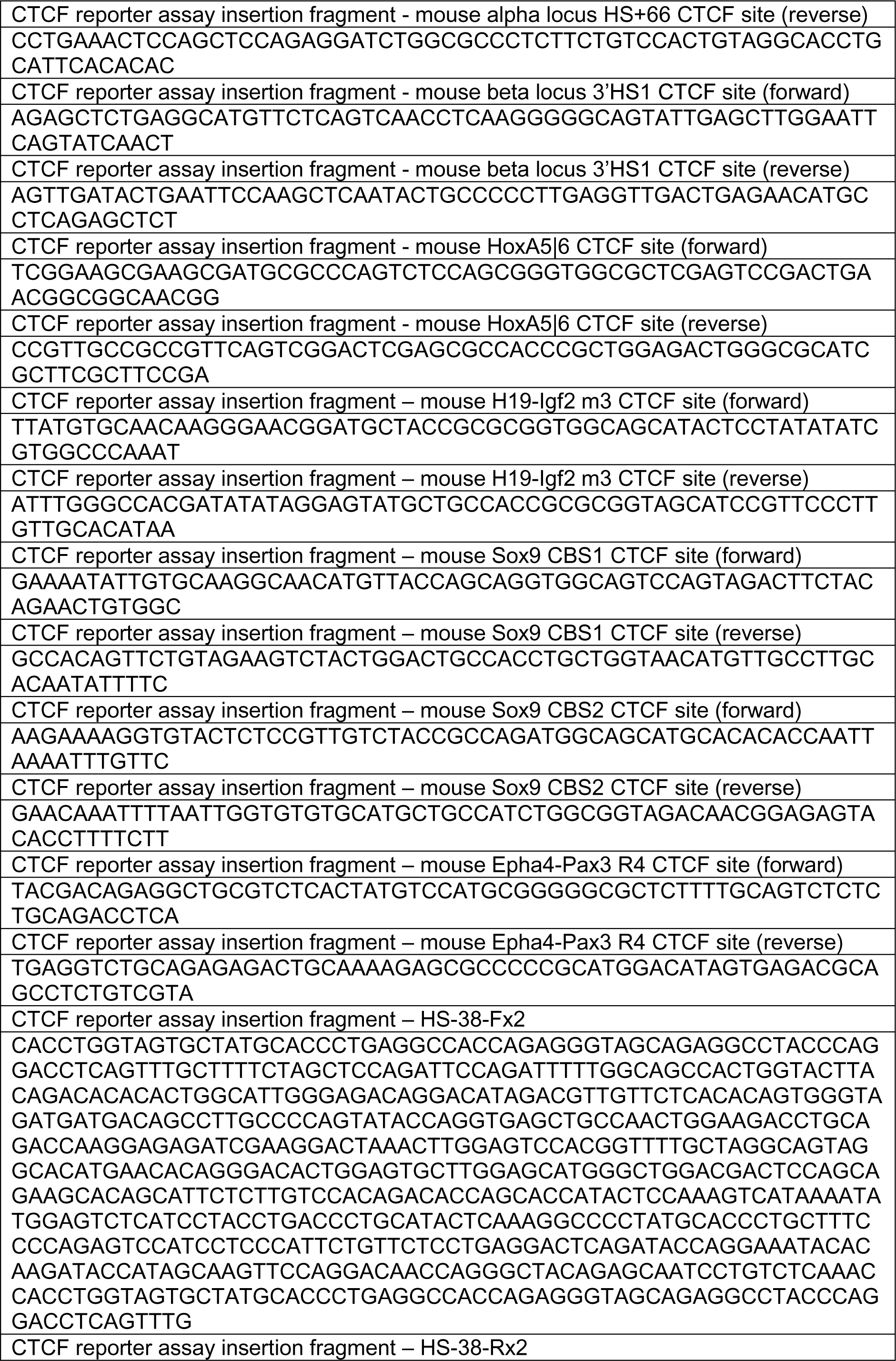
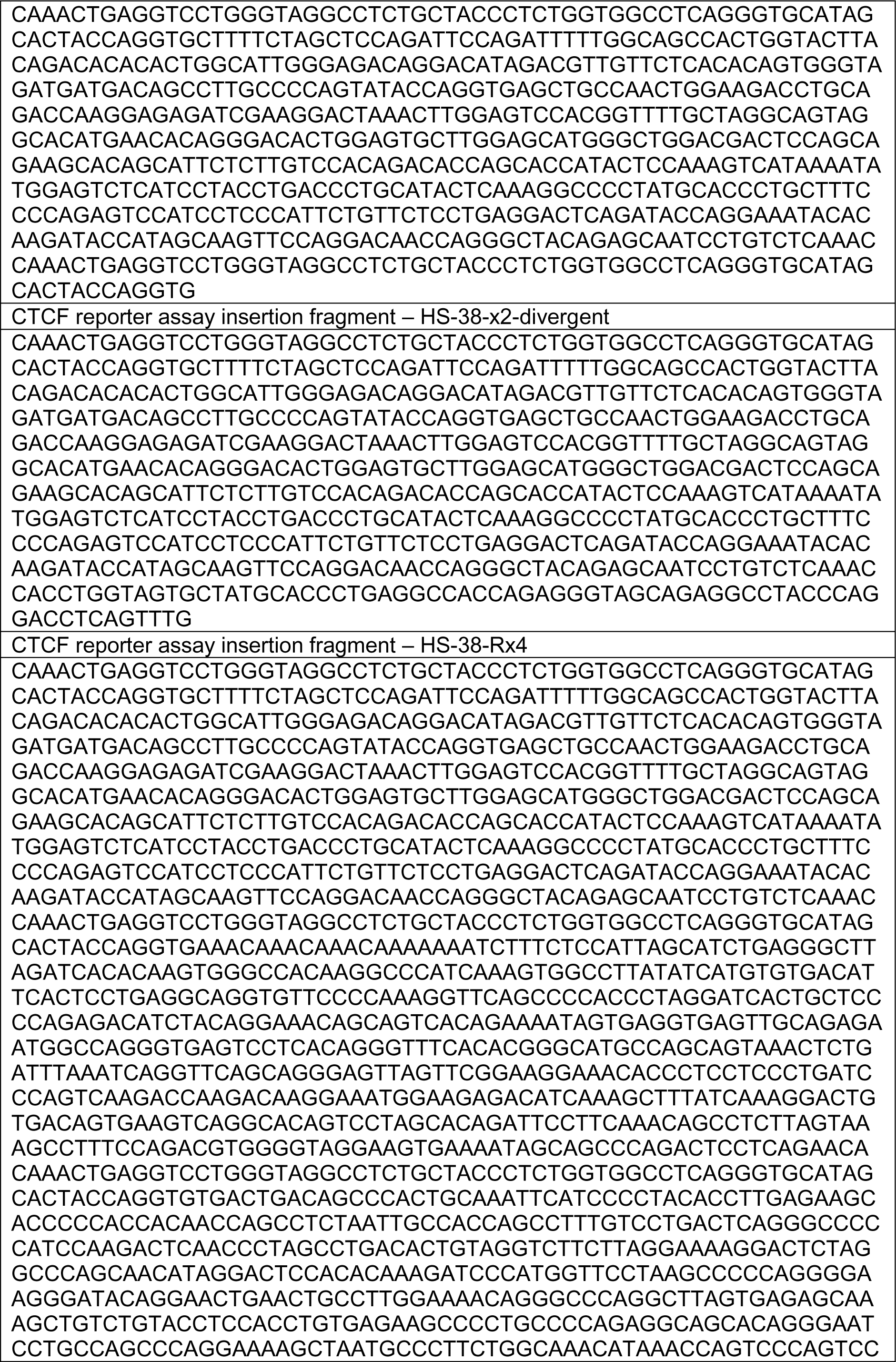
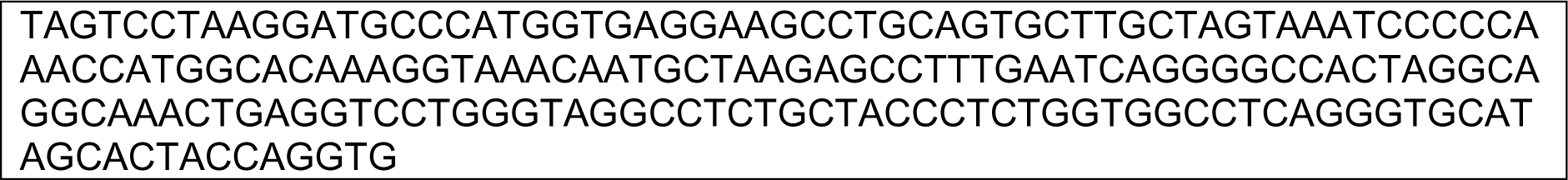
Donor sequences for CRISPR-cas9 editing.

## Data availability

All sequencing data have been submitted to the NCBI Gene Expression Omnibus under accession number GSE163012. Wild-type CTCF ChIP-seq and ATAC-seq data in EBs have previously been published(Francis, Harold et al. 2022), as was wild-type Rad21 ChIP-seq data in spleen(Hanssen, Kassouf et al. 2017).

## Acknowledgments

We would like to thank Jacqueline Sloane-Stanley, Jacqueline Sharpe, Sue Butler, Chris Preece and Daniel Biggs for help with transgenic animals. We thank Jelena Telenius for help with bioinformatic analysis, and Kevin Clark and Sally-Ann Clark of the WIMM Flow Cytometry Facility for help with FACS analysis. This work was supported by Wellcome (Chromosome and Developmental Biology PhD programme, reference 215111/Z/18/Z, to R.J.S., and reference 099684/Z/12/Z, to L.L.P.H.; Core Award Grant reference 203141/Z/16/Z, to B.D.); the Medical Research Council (MRC Core Funding and Centenary Award reference 4050189188, to D.R.H.); and the Chinese Academy of Medical Sciences (CAMS) Innovation Fund for Medical Science (CIFMS), China (grant number: 2018-I2M-2-002, to F.H.T. and D.R.H.).

## Author contributions

L.L.P.H., D.R.H., and B.D. conceived the project. R.J.S, F.H.T, L.L.P.H, M.T.K and B.D. designed experiments. R.J.S., F.H.T., D.J.D., and C.L.H. performed experiments. R.J.S. and D.J.D. performed bioinformatics analyses. R.J.S and F.H.T. performed data analyses. E.G. and J.R.H. designed and generated dTAG cells. R.J.S. and D.R.H. wrote the manuscript. F.H.T. and M.T.K. provided comments on the manuscript. J.R.H., R.A.B, B.D., M.T.K. and D.R.H. supervised the project.

## Notes

### Competing Interest Statement

J.R.H is a founder and shareholder of Nucleome Therapeutics. The remaining authors declare no competing interests.

## References

Aljahani, A., P. Hua, M. A. Karpinska, K. Quililan, J. O. J. Davies and A. M. Oudelaar (2022). “Analysis of sub-kilobase chromatin topology reveals nano-scale regulatory interactions with variable dependence on cohesin and CTCF.” Nat Commun 13(1): 2139.

Banigan, E. J., W. Tang, A. A. van den Berg, R. R. Stocsits, G. Wutz, H. B. Brandão, G. A. Busslinger, J.-M. Peters and L. A. Mirny (2023). “Transcription shapes 3D chromatin organization by interacting with loop extrusion.” Proceedings of the National Academy of Sciences 120(11): e2210480120.

Blackwood, E. M. and J. T. Kadonaga (1998). “Going the distance: a current view of enhancer action.” Science 281(5373): 60–63.

Brown, J. M., N. A. Roberts, B. J. Graham, D. Waithe, C. Lagerholm, J. M. Telenius, S. De Ornellas, A. M. Oudelaar, C. Scott, I. Szczerbal, C. Babbs, M. T. Kassouf, J. R. Hughes, D. R. Higgs and V. J. Buckle (2018). “A tissue-specific self-interacting chromatin domain forms independently of enhancer-promoter interactions.” Nature Communications 9(1): 3849.

Buenrostro, J. D., P. G. Giresi, L. C. Zaba, H. Y. Chang and W. J. Greenleaf (2013). “Transposition of native chromatin for fast and sensitive epigenomic profiling of open chromatin, DNA-binding proteins and nucleosome position.” Nature Methods 10(12): 1213–1218.

Calderon, L., F. D. Weiss, J. A. Beagan, M. S. Oliveira, R. Georgieva, Y.-F. Wang, T. S. Carroll, G. Dharmalingam, W. Gong, K. Tossell, V. de Paola, C. Whilding, M. A. Ungless, A. G. Fisher, J. E. Phillips-Cremins and M. Merkenschlager (2022). “Cohesin-dependence of neuronal gene expression relates to chromatin loop length.” eLife 11: e76539.

Catarino, R. R. and A. Stark (2018). “Assessing sufficiency and necessity of enhancer activities for gene expression and the mechanisms of transcription activation.” Genes Dev 32(3-4): 202–223.

Chen, K., J. Liu, S. Heck, J. A. Chasis, X. An and N. Mohandas (2009). “Resolving the distinct stages in erythroid differentiation based on dynamic changes in membrane protein expression during erythropoiesis.” Proc Natl Acad Sci U S A 106(41): 17413–17418.

Cho, S. W., J. Xu, R. Sun, M. R. Mumbach, A. C. Carter, Y. G. Chen, K. E. Yost, J. Kim, J. He, S. A. Nevins, S. F. Chin, C. Caldas, S. J. Liu, M. A. Horlbeck, D. A. Lim, J. S. Weissman, C. Curtis and H. Y. Chang (2018). “Promoter of lncRNA Gene PVT1 Is a Tumor-Suppressor DNA Boundary Element.” Cell 173(6): 1398–1412 e1322.

Cong, L., F. A. Ran, D. Cox, S. Lin, R. Barretto, N. Habib, P. D. Hsu, X. Wu, W. Jiang, L. A. Marraffini and F. Zhang (2013). “Multiplex Genome Engineering Using CRISPR/Cas Systems.” Science 339(6121): 819–823.

Cuartero, S., F. D. Weiss, G. Dharmalingam, Y. Guo, E. Ing-Simmons, S. Masella, I. Robles-Rebollo, X. Xiao, Y.-F. Wang, I. Barozzi, D. Djeghloul, M. T. Amano, H. Niskanen, E. Petretto, R. D. Dowell, K. Tachibana, M. U. Kaikkonen, K. A. Nasmyth, B. Lenhard, G. Natoli, A. G. Fisher and M. Merkenschlager (2018). “Control of inducible gene expression links cohesin to hematopoietic progenitor self-renewal and differentiation.” Nature Immunology 19(9): 932–941.

Davidson, I. F., R. Barth, M. Zaczek, J. van der Torre, W. Tang, K. Nagasaka, R. Janissen, J. Kerssemakers, G. Wutz, C. Dekker and J. M. Peters (2023). “CTCF is a DNA-tension-dependent barrier to cohesin-mediated loop extrusion.” Nature 616(7958): 822–827.

Davies, J. O., A. M. Oudelaar, D. R. Higgs and J. R. Hughes (2017). “How best to identify chromosomal interactions: a comparison of approaches.” Nat Methods 14(2): 125–134.

Davies, J. O. J., J. M. Telenius, S. J. McGowan, N. A. Roberts, S. Taylor, D. R. Higgs and J. R. Hughes (2016). “Multiplexed analysis of chromosome conformation at vastly improved sensitivity.” Nature methods 13(1): 74–80.

de Wit, E. and E. P. Nora (2023). “New insights into genome folding by loop extrusion from inducible degron technologies.” Nat Rev Genet 24(2): 73–85.

De Wit, E., E. S. Vos, S. J. Holwerda, C. Valdes-Quezada, M. J. Verstegen, H. Teunissen, E. Splinter, P. J. Wijchers, P. H. Krijger and W. de Laat (2015). “CTCF Binding Polarity Determines Chromatin Looping.” Molecular Cell 60(4): 676–684.

Deng, W., J. Lee, H. Wang, J. Miller, A. Reik, Philip D. Gregory, A. Dean and Gerd A. Blobel (2012). “Controlling Long-Range Genomic Interactions at a Native Locus by Targeted Tethering of a Looping Factor.” Cell 149(6): 1233–1244.

Dixon, J. R., S. Selvaraj, F. Yue, A. Kim, Y. Li, Y. Shen, M. Hu, J. S. Liu and B. Ren (2012). “Topological domains in mammalian genomes identified by analysis of chromatin interactions.” Nature 485(7398): 376–380.

Downes, D. J., A. L. Smith, M. A. Karpinska, T. Velychko, K. Rue-Albrecht, D. Sims, T. A. Milne, J. O. J. Davies, A. M. Oudelaar and J. R. Hughes (2022). “Capture-C: a modular and flexible approach for high-resolution chromosome conformation capture.” Nat Protoc 17(2): 445–475.

Engel, J. D. and K. Tanimoto (2000). “Looping, Linking, and Chromatin Activity: New Insights into &#x3b2;-*globin* Locus Regulation.” Cell 100(5): 499–502.

Francis, H. S., C. L. Harold, R. A. Beagrie, A. J. King, M. E. Gosden, J. W. Blayney, D. M. Jeziorska, C. Babbs, D. R. Higgs and M. T. Kassouf (2022). “Scalable in vitro production of defined mouse erythroblasts.” PLoS One 17(1): e0261950.

Fudenberg, G., N. Abdennur, M. Imakaev, A. Goloborodko and L. A. Mirny (2017). “Emerging Evidence of Chromosome Folding by Loop Extrusion.” Cold Spring Harb Symp Quant Biol 82: 45–55.

Fudenberg, G., M. Imakaev, C. Lu, A. Goloborodko, N. Abdennur and Leonid A. Mirny (2016). “Formation of Chromosomal Domains by Loop Extrusion.” Cell Reports 15(9): 2038–2049.

Furlong, E. E. M. and M. Levine (2018). “Developmental enhancers and chromosome topology.” Science 361(6409): 1341–1345.

Goel, V. Y., M. K. Huseyin and A. S. Hansen (2023). “Region Capture Micro-C reveals coalescence of enhancers and promoters into nested microcompartments.” Nat Genet 55(6): 1048–1056.

Grosveld, F., J. van Staalduinen and R. Stadhouders (2021). “Transcriptional Regulation by (Super)Enhancers: From Discovery to Mechanisms.” Annu Rev Genomics Hum Genet 22: 127–146.

Guo, Y., Q. Xu, D. Canzio, J. Shou, J. Li, David U. Gorkin, I. Jung, H. Wu, Y. Zhai, Y. Tang, Y. Lu, Y. Wu, Z. Jia, W. Li, Michael Q. Zhang, B. Ren, Adrian R. Krainer, T. Maniatis and Q. Wu (2015). “CRISPR Inversion of CTCF Sites Alters Genome Topology and Enhancer/Promoter Function.” Cell 162(4): 900–910.

Hanssen, L. L. P., M. T. Kassouf, A. M. Oudelaar, D. Biggs, C. Preece, D. J. Downes, M. Gosden, J. A. Sharpe, J. A. Sloane-Stanley, J. R. Hughes, B. Davies and D. R. Higgs (2017). “Tissue-specific CTCF–cohesin-mediated chromatin architecture delimits enhancer interactions and function in vivo.” Nature Cell Biology 19(8): 952–961.

Harrold, C. L., M. E. Gosden, L. L. P. Hanssen, R. J. Stolper, D. J. Downes, J. M. Telenius, D. Biggs, C. Preece, S. Alghadban, J. A. Sharpe, B. Davies, J. A. Sloane-Stanley, M. T. Kassouf, J. R. Hughes and D. R. Higgs (2020). “A functional overlap between actively transcribed genes and chromatin boundary elements.” bioRxiv: 2020.2007.2001.182089.

Hay, D., J. R. Hughes, C. Babbs, J. O. J. Davies, B. J. Graham, L. Hanssen, M. T. Kassouf, A. M. Marieke Oudelaar, J. A. Sharpe, M. C. Suciu, J. Telenius, R. Williams, C. Rode, P.-S. Li, L. A. Pennacchio, J. A. Sloane-Stanley, H. Ayyub, S. Butler, T. Sauka-Spengler, R. J. Gibbons, A. J. H. Smith, W. G. Wood and D. R. Higgs (2016). “Genetic dissection of the α- globin super-enhancer in vivo.” Nature genetics 48(8): 895–903.

Hentges, L. D., M. J. Sergeant, D. J. Downes, J. R. Hughes and S. Taylor (2021). “LanceOtron: a deep learning peak caller for ATAC-seq, ChIP-seq, and DNase-seq.” biorxiv.

Hsieh, T. S., C. Cattoglio, E. Slobodyanyuk, A. S. Hansen, X. Darzacq and R. Tjian (2022). “Enhancer-promoter interactions and transcription are largely maintained upon acute loss of CTCF, cohesin, WAPL or YY1.” Nat Genet.

Hua, P., M. Badat, L. L. P. Hanssen, L. D. Hentges, N. Crump, D. J. Downes, D. M. Jeziorska, A. M. Oudelaar, R. Schwessinger, S. Taylor, T. A. Milne, J. R. Hughes, D. R. Higgs and J. O. J. Davies (2021). “Defining genome architecture at base-pair resolution.” Nature 595(7865): 125–129.

Huang, H., Q. Zhu, A. Jussila, Y. Han, B. Bintu, C. Kern, M. Conte, Y. Zhang, S. Bianco, A. M. Chiariello, M. Yu, R. Hu, M. Tastemel, I. Juric, M. Hu, M. Nicodemi, X. Zhuang and B. Ren (2021). “CTCF mediates dosage- and sequence-context-dependent transcriptional insulation by forming local chromatin domains.” Nat Genet.

Jackson, D. A., F. J. Iborra, E. M. M. Manders and P. R. Cook (1998). “Numbers and Organization of RNA Polymerases, Nascent Transcripts, and Transcription Units in HeLa Nuclei.” Molecular Biology of the Cell 9(6): 1523–1536.

Kagey, M. H., J. J. Newman, S. Bilodeau, Y. Zhan, D. A. Orlando, N. L. van Berkum, C. C. Ebmeier, J. Goossens, P. B. Rahl, S. S. Levine, D. J. Taatjes, J. Dekker and R. A. Young (2010). “Mediator and cohesin connect gene expression and chromatin architecture.” Nature 467(7314): 430–435.

Keller, G. M. (1995). “In vitro differentiation of embryonic stem cells.” Curr Opin Cell Biol 7(6): 862–869.

Kent, W. J., C. W. Sugnet, T. S. Furey, K. M. Roskin, T. H. Pringle, A. M. Zahler, Haussler and David (2002). “The Human Genome Browser at UCSC.” Genome Research 12(6): 996–1006.

Khoury, A., J. Achinger-Kawecka, S. A. Bert, G. C. Smith, H. J. French, P. L. Luu, T. J. Peters, Q. Du, A. J. Parry, F. Valdes-Mora, P. C. Taberlay, C. Stirzaker, A. L. Statham and S. J. Clark (2020). “Constitutively bound CTCF sites maintain 3D chromatin architecture and long-range epigenetically regulated domains.” Nat Commun 11(1): 54.

King, A. J., D. Songdej, D. J. Downes, R. A. Beagrie, S. Liu, M. Buckley, P. Hua, M. C. Suciu, A. Marieke Oudelaar, L. L. P. Hanssen, D. Jeziorska, N. Roberts, S. J. Carpenter, H. Francis, J. Telenius, A. A. Olijnik, J. A. Sharpe, J. Sloane-Stanley, J. Eglinton, M. T. Kassouf, S. H. Orkin, L. A. Pennacchio, J. O. J. Davies, J. R. Hughes, D. R. Higgs and C. Babbs (2021). “Reactivation of a developmentally silenced embryonic globin gene.” Nat Commun 12(1): 4439.

Langmead, B. and S. L. Salzberg (2012). “Fast gapped-read alignment with Bowtie 2.” Nature Methods 9(4): 357–359.

Lee, G. and I. Saito (1998). “Role of nucleotide sequences of loxP spacer region in Cre-mediated recombination.” Gene 216(1): 55–65.

Li, H., B. Handsaker, A. Wysoker, T. Fennell, J. Ruan, N. Homer, G. Marth, G. Abecasis and R. Durbin (2009). “The Sequence Alignment/Map format and SAMtools.” Bioinformatics 25(16): 2078–2079.

Li, Y., J. H. I. Haarhuis, Á. Sedeño Cacciatore, R. Oldenkamp, M. S. van Ruiten, L. Willems, H. Teunissen, K. W. Muir, E. de Wit, B. D. Rowland and D. Panne (2020). “The structural basis for cohesin–CTCF-anchored loops.” Nature 578(7795): 472–476.

Liu, N. Q., M. Maresca, T. van den Brand, L. Braccioli, M. Schijns, H. Teunissen, B. G. Bruneau, E. P. Nora and E. de Wit (2021). “WAPL maintains a cohesin loading cycle to preserve cell-type-specific distal gene regulation.” Nat Genet 53(1): 100–109.

Long, H. K., S. L. Prescott and J. Wysocka (2016). “Ever-Changing Landscapes: Transcriptional Enhancers in Development and Evolution.” Cell 167(5): 1170–1187.

Lupianez, D. G., K. Kraft, V. Heinrich, P. Krawitz, F. Brancati, E. Klopocki, D. Horn, H. Kayserili, J. M. Opitz, R. Laxova, F. Santos-Simarro, B. Gilbert-Dussardier, L. Wittler, M. Borschiwer, S. A. Haas, M. Osterwalder, M. Franke, B. Timmermann, J. Hecht, M. Spielmann, A. Visel and S. Mundlos (2015). “Disruptions of topological chromatin domains cause pathogenic rewiring of gene-enhancer interactions.” Cell 161(5): 1012–1025.

Lupianez, D. G., M. Spielmann and S. Mundlos (2016). “Breaking TADs: How Alterations of Chromatin Domains Result in Disease.” Trends Genet 32(4): 225–237.

Luppino, J. M., D. S. Park, S. C. Nguyen, Y. Lan, Z. Xu, R. Yunker and E. F. Joyce (2020). “Cohesin promotes stochastic domain intermingling to ensure proper regulation of boundary-proximal genes.” Nature Genetics.

Mao, X., Y. Fujiwara and S. H. Orkin (1999). “Improved reporter strain for monitoring Cre recombinase-mediated DNA excisions in mice.” Proceedings of the National Academy of Sciences 96(9): 5037–5042.

Martinez-Ara, M., F. Comoglio, J. van Arensbergen and B. van Steensel (2022). “Systematic analysis of intrinsic enhancer-promoter compatibility in the mouse genome.” Mol Cell 82(13): 2519–2531 e2516.

Mattingly, M., C. Seidel, S. Munoz, Y. Hao, Y. Zhang, Z. Wen, L. Florens, F. Uhlmann and J. L. Gerton (2022). “Mediator recruits the cohesin loader Scc2 to RNA Pol II-transcribed genes and promotes sister chromatid cohesion.” Curr Biol 32(13): 2884–2896 e2886.

Nabet, B., J. M. Roberts, D. L. Buckley, J. Paulk, S. Dastjerdi, A. Yang, A. L. Leggett, M. A. Erb, M. A. Lawlor, A. Souza, T. G. Scott, S. Vittori, J. A. Perry, J. Qi, G. E. Winter, K. K. Wong, N. S. Gray and J. E. Bradner (2018). “The dTAG system for immediate and target-specific protein degradation.” Nat Chem Biol 14(5): 431–441.

Nakahashi, H., K.-Rim K. Kwon, W. Resch, L. Vian, M. Dose, D. Stavreva, O. Hakim, N. Pruett, S. Nelson, A. Yamane, J. Qian, W. Dubois, S. Welsh, Robert D. Phair, B. F. Pugh, V. Lobanenkov, Gordon L. Hager and R. Casellas (2013). “A Genome-wide Map of CTCF Multivalency Redefines the CTCF Code.” Cell Reports 3(5): 1678–1689.

Nora, E. P., A. Goloborodko, A. L. Valton, J. H. Gibcus, A. Uebersohn, N. Abdennur, J. Dekker, L. A. Mirny and B. G. Bruneau (2017). “Targeted Degradation of CTCF Decouples Local Insulation of Chromosome Domains from Genomic Compartmentalization.” Cell 169(5): 930–944.e922.

Nora, E. P., B. R. Lajoie, E. G. Schulz, L. Giorgetti, I. Okamoto, N. Servant, T. Piolot, N. L. van Berkum, J. Meisig, J. Sedat, J. Gribnau, E. Barillot, N. Bluthgen, J. Dekker and E. Heard (2012). “Spatial partitioning of the regulatory landscape of the X-inactivation centre.” Nature 485(7398): 381–385.

Oudelaar, A. M., R. A. Beagrie, M. Gosden, S. de Ornellas, E. Georgiades, J. Kerry, D. Hidalgo, J. Carrelha, A. Shivalingam, A. H. El-Sagheer, J. M. Telenius, T. Brown, V. J. Buckle, M. Socolovsky, D. R. Higgs and J. R. Hughes (2020). “Dynamics of the 4D genome during in vivo lineage specification and differentiation.” Nat Commun 11(1): 2722.

Pugacheva, E. M., N. Kubo, D. Loukinov, M. Tajmul, S. Kang, A. L. Kovalchuk, A. V. Strunnikov, G. E. Zentner, B. Ren and V. V. Lobanenkov (2020). “CTCF mediates chromatin looping via N-terminal domain-dependent cohesin retention.” Proceedings of the National Academy of Sciences 117(4): 2020–2031.

Quinlan, A. R. and I. M. Hall (2010). “BEDTools: a flexible suite of utilities for comparing genomic features.” Bioinformatics 26(6): 841–842.

Ramírez, F., D. P. Ryan, B. Grüning, V. Bhardwaj, F. Kilpert, A. S. Richter, S. Heyne, F. Dündar and T. Manke (2016). “deepTools2: a next generation web server for deep-sequencing data analysis.” Nucleic Acids Res 44(W1): W160–165.

Ran, F. A., P. D. Hsu, J. Wright, V. Agarwala, D. A. Scott and F. Zhang (2013). “Genome engineering using the CRISPR-Cas9 system.” Nature Protocols 8(11): 2281–2308.

Rao, S. S., M. H. Huntley, N. C. Durand, E. K. Stamenova, I. D. Bochkov, J. T. Robinson, A. L. Sanborn, I. Machol, A. D. Omer, E. S. Lander and E. L. Aiden (2014). “A 3D map of the human genome at kilobase resolution reveals principles of chromatin looping.” Cell 159(7): 1665–1680.

Rao, S. S. P., S.-C. Huang, B. Glenn St Hilaire, J. M. Engreitz, E. M. Perez, K.-R. Kieffer-Kwon, A. L. Sanborn, S. E. Johnstone, G. D. Bascom, I. D. Bochkov, X. Huang, M. S. Shamim, J. Shin, D. Turner, Z. Ye, A. D. Omer, J. T. Robinson, T. Schlick, B. E. Bernstein, R. Casellas, E. S. Lander and E. L. Aiden (2017). “Cohesin Loss Eliminates All Loop Domains.” Cell 171(2): 305–320.e324.

Rinzema, N. J., K. Sofiadis, S. J. D. Tjalsma, M. Verstegen, Y. Oz, C. Valdes-Quezada, A. K. Felder, T. Filipovska, S. van der Elst, Z. de Andrade Dos Ramos, R. Han, P. H. L. Krijger and W. de Laat (2022). “Building regulatory landscapes reveals that an enhancer can recruit cohesin to create contact domains, engage CTCF sites and activate distant genes.” Nat Struct Mol Biol 29(6): 563–574.

Robles-Rebollo, I., S. Cuartero, A. Canellas-Socias, S. Wells, M. M. Karimi, E. Mereu, A. G. Chivu, H. Heyn, C. Whilding, D. Dormann, S. Marguerat, I. Rioja, R. K. Prinjha, M. P. H. Stumpf, A. G. Fisher and M. Merkenschlager (2022). “Cohesin couples transcriptional bursting probabilities of inducible enhancers and promoters.” Nat Commun 13(1): 4342.

Sanborn, A. L., S. S. P. Rao, S.-C. Huang, N. C. Durand, M. H. Huntley, A. I. Jewett, I. D. Bochkov, D. Chinnappan, A. Cutkosky, J. Li, K. P. Geeting, A. Gnirke, A. Melnikov, D. McKenna, E. K. Stamenova, E. S. Lander and E. L. Aiden (2015). “Chromatin extrusion explains key features of loop and domain formation in wild-type and engineered genomes.” Proceedings of the National Academy of Sciences 112(47): E6456–E6465.

Schwarzer, W., N. Abdennur, A. Goloborodko, A. Pekowska, G. Fudenberg, Y. Loe-Mie, N. A. Fonseca, W. Huber, C. H. Haering and L. Mirny (2017). “Two independent modes of chromatin organization revealed by cohesin removal.” Nature 551(7678): 51–56.

Spivak, J. L., D. Toretti and H. W. Dickerman (1973). “Effect of Phenylhydrazine-induced Hemolytic Anemia on Nuclear RNA Polymerase Activity of the Mouse Spleen.” Blood 42(2): 257–266.

Symmons, O., V. V. Uslu, T. Tsujimura, S. Ruf, S. Nassari, W. Schwarzer, L. Ettwiller and F. Spitz (2014). “Functional and topological characteristics of mammalian regulatory domains.” Genome Research 24(3): 390–400.

Taniguchi, M., M. Sanbo, S. Watanabe, I. Naruse, M. Mishina and T. Yagi (1998). “Efficient production of Cre-mediated site-directed recombinants through the utilization of the puromycin resistance gene, pac: A transient gene-integration marker for ES cells.” Nucleic Acids Research 26(2): 679–680.

Telenius, J. M., D. J. Downes, M. Sergeant, A. M. Oudelaar, S. McGowan, J. Kerry, L. L. P. Hanssen, R. Schwessinger, C. Q. Eijsbouts, J. O. J. Davies, S. Taylor and J. R. Hughes (2020). “CaptureCompendium: a comprehensive toolkit for 3C analysis.” biorxiv: 2020.2002.2017.952572.

Thiecke, M. J., G. Wutz, M. Muhar, W. Tang, S. Bevan, V. Malysheva, R. Stocsits, T. Neumann, J. Zuber, P. Fraser, S. Schoenfelder, J. M. Peters and M. Spivakov (2020). “Cohesin-Dependent and -Independent Mechanisms Mediate Chromosomal Contacts between Promoters and Enhancers.” Cell Reports 32(3): 107929.

Tsang, F. H., R. J. Stolper, H. Muhammad, L. J. Cornell, B. Davies, M. T. Kassouf and D. R. Higgs (2023). “The characteristics of CTCF binding sequences contribute to enhancer blocking activity “ Biorxiv.

Tsujimura, T., F. A. Klein, K. Langenfeld, J. Glaser, W. Huber and F. Spitz (2015). “A Discrete Transition Zone Organizes the Topological and Regulatory Autonomy of the Adjacent Tfap2c and Bmp7 Genes.” PLOS Genetics 11(1): e1004897.

von Lindern, M., E. M. Deiner, H. Dolznig, M. Parren-Van Amelsvoort, M. J. Hayman, E. W. Mullner and H. Beug (2001). “Leukemic transformation of normal murine erythroid progenitors: v- and c-ErbB act through signaling pathways activated by the EpoR and c-Kit in stress erythropoiesis.” Oncogene 20(28): 3651–3664.

Vos, E. S. M., C. Valdes-Quezada, Y. Huang, A. Allahyar, M. Verstegen, A. K. Felder, F. van der Vegt, E. C. H. Uijttewaal, P. H. L. Krijger and W. de Laat (2021). “Interplay between CTCF boundaries and a super enhancer controls cohesin extrusion trajectories and gene expression.” Molecular Cell.

Wutz, G., C. Várnai, K. Nagasaka, D. A. Cisneros, R. R. Stocsits, W. Tang, S. Schoenfelder, G. Jessberger, M. Muhar and M. J. Hossain (2017). “Topologically associating domains and chromatin loops depend on cohesin and are regulated by CTCF, WAPL, and PDS5 proteins.” The EMBO journal 36(24): 3573–3599.

Zabidi, M. A., C. D. Arnold, K. Schernhuber, M. Pagani, M. Rath, O. Frank and A. Stark (2015). “Enhancer-core-promoter specificity separates developmental and housekeeping gene regulation.” Nature 518(7540): 556–559.

Zuin, J., G. Roth, Y. Zhan, J. Cramard, J. Redolfi, E. Piskadlo, P. Mach, M. Kryzhanovska, G. Tihanyi, H. Kohler, M. Eder, C. Leemans, B. van Steensel, P. Meister, S. Smallwood and L. Giorgetti (2022). “Nonlinear control of transcription through enhancer-promoter interactions.” Nature 604(7906): 571–577.

