## Supplementary figures for "Loop extrusion by cohesin plays a key role in enhancer-activated gene expression during differentiation"

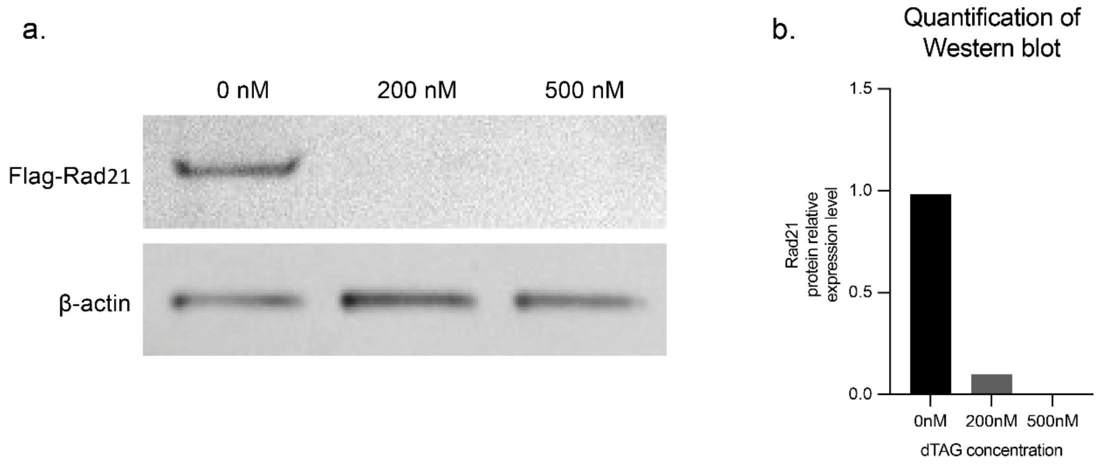


*Supplementary figure 1. Western blot analysis of protein extract from differentiated Rad21-FKBP12$^{F36V}$ mESCs. a) Western blot of protein extract of CD71+ erythroid cells treated with varying concentrations of dTAG. The Rad21 fusion protein was detected with a FLAG antibody and beta-actin was used as a loading control. b) Quantification of Rad21 proteins levels after dTAG treatment as determined by western blot.*

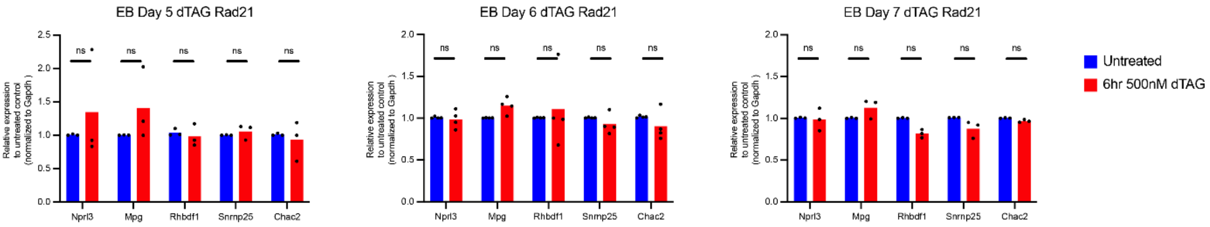


*Supplementary figure 2. Gene expression changes upon Rad21 depletion in non-erythroid genes. Gene expression was measured by qPCR and normalised to Gapdh in day 5, day 6 and day 7 EBs with or without dTAG treatment. P-values were obtained by an unpaired two-tailed Student's t-test: Not significant (NS) p > 0.05.*

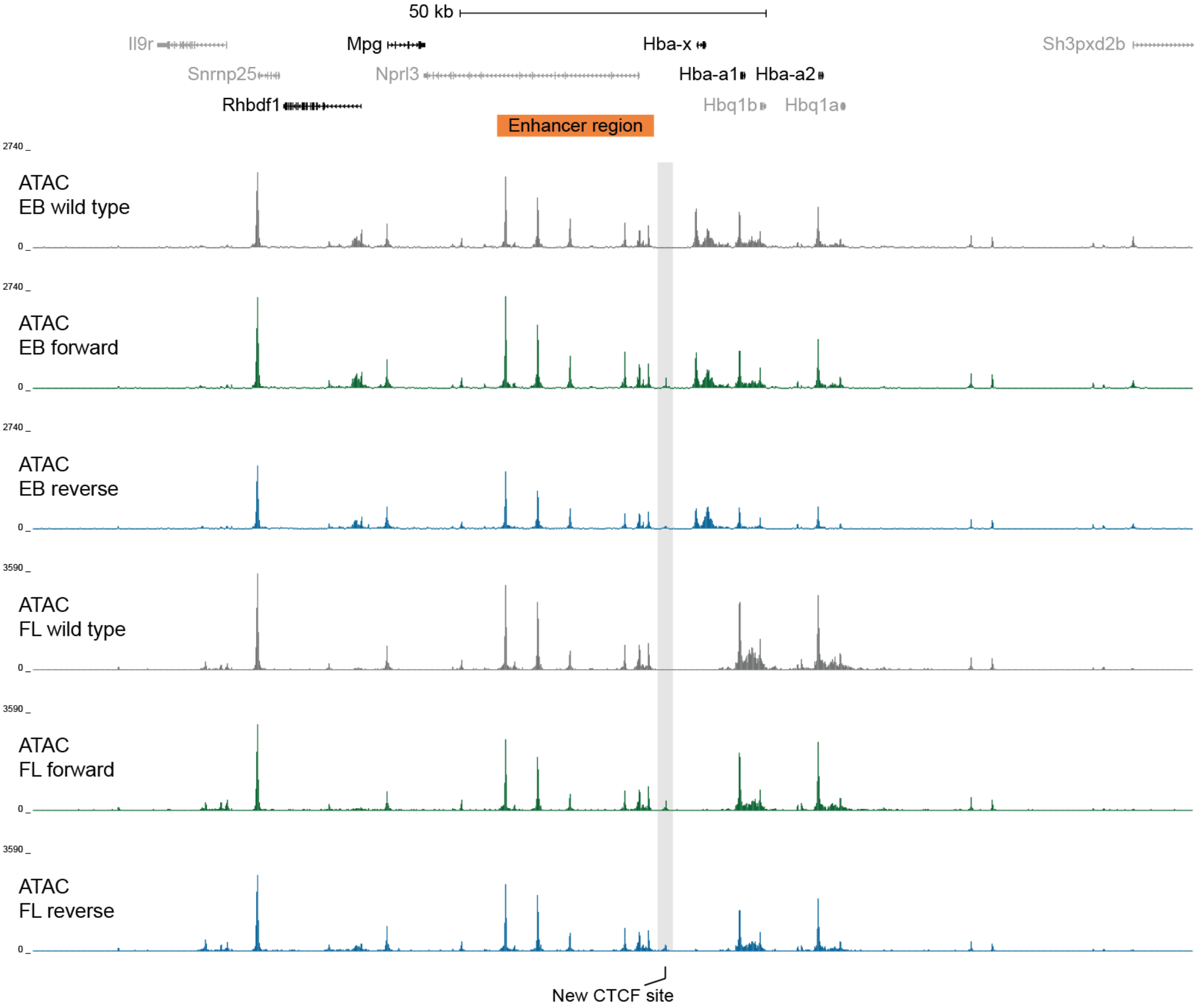


*Supplementary figure 3. ATAC-seq tracks in the Forward and Reverse model. RPKM-normalised ATAC-seq tracks in WT (grey), Forward (green), and Reverse (blue) erythroid cells either from day 7 EBs or E12.5 fetal liver. Three replicates were merged for each track.*

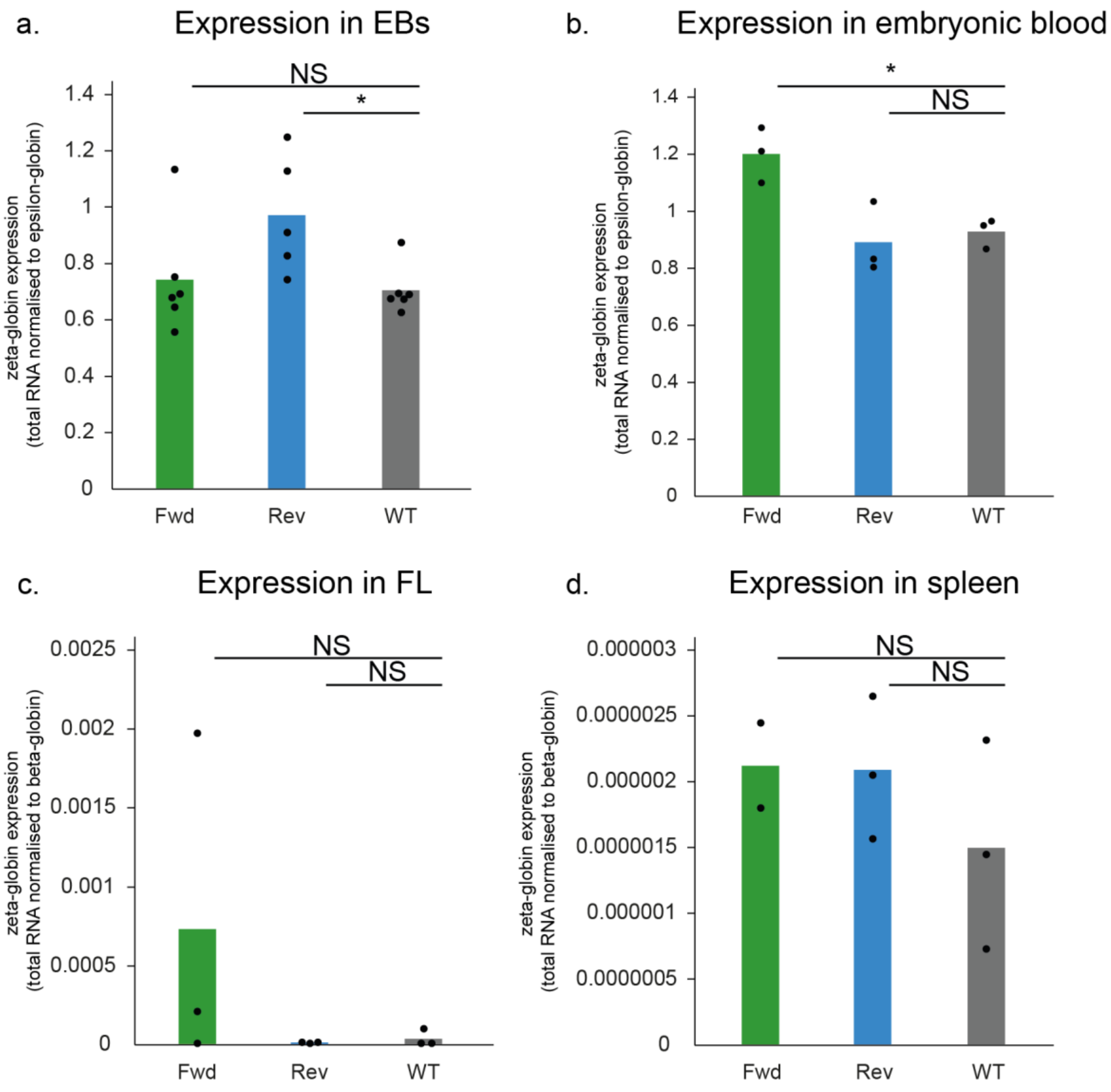


*Supplementary figure 4. Expression of zeta-globin in various erythroid tissues in the Forward and Reverse models. a) Expression of the embryonic zeta-globin (Hba-x) in EB-derived CD71+ erythroid cells, normalised to the embryonic beta-like globin, epsilon-globin (Hbb-y; n=5 for Rev, n=6 for Fwd and WT). Bar plot shows the mean expression with individual data points marked by black dots. b) Expression of zeta-globin in E10.5 embryonic blood normalised to epsilon-globin (n=3 for all). c) Expression of zeta-globin normalised to beta-globin in cultured E12.5 fetal liver Ter119+ erythroid cells (n=3 for all). d) Expression of zeta-globin normalised to beta-globin in spleen-derived Ter119+ erythroid cells (n=2 for Fwd, n=3 for Rev and WT). P-values were obtained using an unpaired two-tailed Student's t-test. Not significant (NS) p > 0.05, * p <0.05.*

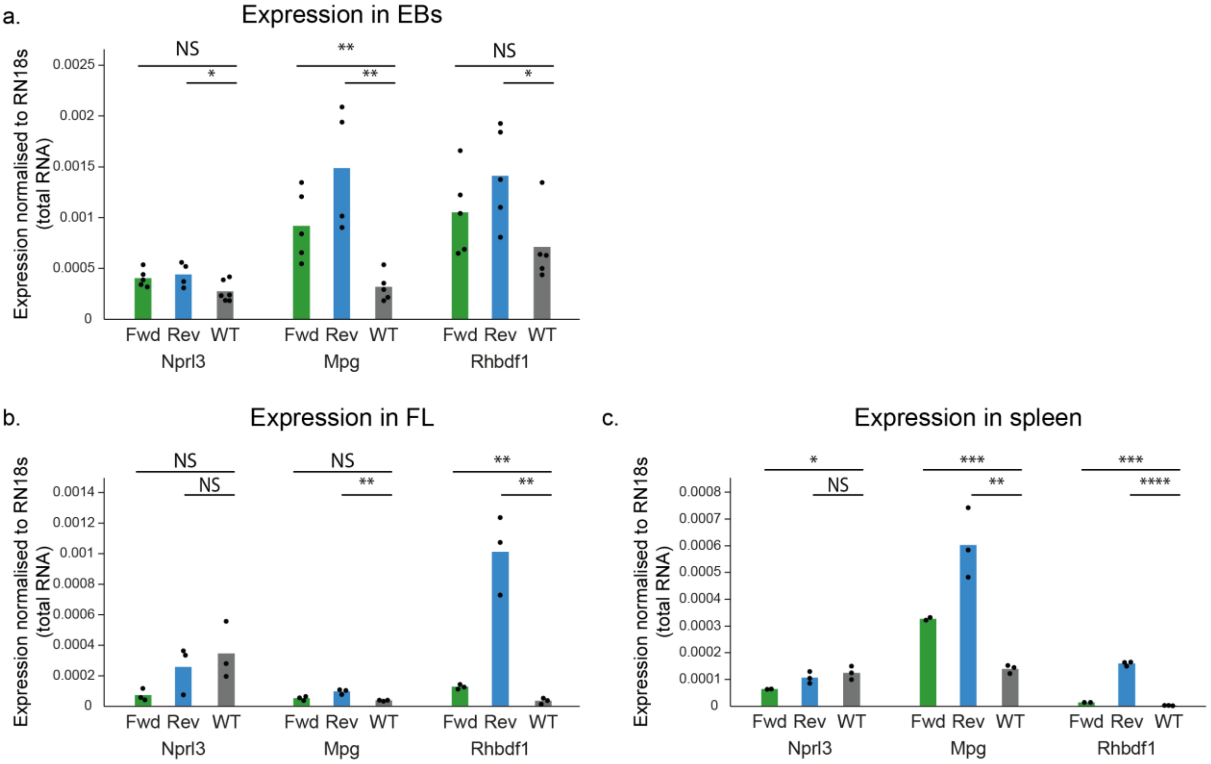


*Supplementary figure 5. Expression of non-erythroid genes at the alpha-globin locus in the Forward and Reverse model. a) Expression of Nprl3 (Fwd n=5, Rev n=4, WT n= 6), Mpg (Fwd and WT n=5, Rev n=4), and Rhbdf1 (n=5 for all) in EB-derived CD71+ erythroid cells, normalised to Rn18s. Bar plot shows the mean expression with individual data points marked by black dots. b) Expression of Nprl3, Mpg and Rhbdf1 in cultured E12.5 fetal liver Ter119+ erythroid cells (n=3 for all), normalised to Rn18s. c) Expression of Nprl3, Mpg and Rhbdf1 in spleen-derived Ter119+ erythroid cells (n=2 for Fwd, n=3 for Rev and WT), normalised to Rn18s. P-values were obtained using an unpaired two-tailed Student's t-test. Not significant (NS) p > 0.05, * p <0.05, ** p < 0.01, *** p <0.001, **** p<0.0001.*
